# Taxonomic-free metagenome GWAS to identify gut microbiome functions influencing host phenotypes

**DOI:** 10.64898/2026.01.15.699651

**Authors:** Raphaël Malak, Arthur Frouin, Léo Henches, Antoine Auvergne, Milieu Intérieur Consortium, Christophe Boetto, Harry Sokol, Rayan Chikhi, Hugues Aschard

## Abstract

Genome-wide association studies (GWAS) have been pivotal for uncovering the genetics of human phenotypes, and there is now growing interest in applying GWAS-like methods to explore the role of the microbiome in human health. Here, we developed a taxonomy free GWAS approach that uses *k*-mers, i.e. DNA words of length *k* that can capture single nucleotide polymorphisms, insertion or deletion events, and gene presence–absence, to interrogate the gut microbiome. We applied this method to 26 traits spanning demographics, physiological measurements, health, and lifestyle in 938 healthy participants from the Milieu Intérieur cohort. We generated a *k*-mer abundance matrix encompassing 97 million distinct *k*-mers. GWAS of the 26 traits identified significant associations for seven of them: age, sex, depression, appetite, cooked meat consumption, soda intake, and smoking. By modeling the correlation structure among *k*-mers, we identified a modest number of independent signals and conducted a comprehensive *in silico* functional annotation of these signals, revealing potential mechanisms of host–microbiota interaction. Overall, our analyses demonstrate that *k*-mers can capture biologically relevant functions shared across multiple taxa and provide a refined modeling framework that complements the standard taxonomic-based screening approach.

## Introduction

The vast majority of metagenomic studies rely on taxonomic profiling of samples. The proportion of bacterial species in a metagenome is first estimated from reference genome databases^1,2^ using tools such as Kraken2^3^ or MetaPhlAn4^4^, after which the inferred taxa are examined as the primary variables of interest^5,6^. This workflow has been especially prevalent in gut-microbiome research, where it has successfully identified associations with a broad spectrum of human phenotypes^7,8^. However, there is increasing evidence that examining additional characteristics of the microbiome can further our understanding of the host–microbiome relationship. Taxa co-abundance studies^9,10^, community-based approaches^9^, and multivariate analyses^11^ have identified mechanisms missed by univariate taxa-based analyses. As for single-bacterium studies^12–14^, interest is growing in investigating the association between host phenotypes and the genetics of the gut microbiome using genome-wide association studies (GWAS)—a methodology that has proved reliable, replicable, and feasible for capturing the genetic component of many complex human phenotypes^15–17^. In this direction, Zahavi *et al.* ^18^ recently implemented a GWAS framework that focuses on single-nucleotide polymorphisms (SNPs) extracted from reference genome databases. Although limited in genetic coverage—using only SNPs from the core genome and discarding other sources of variability such as gene presence–absence and the entire accessory genome—this work demonstrated the potential of conducting sequence-based GWAS of the metagenome. GWAS were also performed on collections of isolate bacterial species using k-mers (DNA words of length *k*^12,19,20^), although not on multiple species jointly^12,21^.

Here we present a novel taxonomic-free, *k*-mer-based GWAS approach that captures genetic variability across the entire metagenome. We applied the method to screen for associations between the gut microbiome and phenotypes in 938 healthy participants from the Milieu Intérieur cohort. Given the extensive DNA-sequence sharing among bacteria—resulting from prokaryote-specific genetic exchange processes that can occur in a metagenome (transformation, transduction, and conjugation)^22,23^— we hypothesize that a *k*-mer-based GWAS may detect host-phenotype associations driven by sequences shared across multiple species. Furthermore, bacterial species that are linked to a host phenotype typically possess a wide array of biological functions; identifying effects at the sequence level may help refining the molecular mechanisms underlying those associations. In practice, we first assess and mitigate the computational burden of enumerating and handling billions of *k*-mers derived from tens of billions of reads across hundreds of individuals. When applied to the Milieu Intérieur data, this pipeline yields a unique overview of the cross-taxonomic distribution of species and *k*-mers. We then developed a computationally efficient tool, MOGS (Metagenome Occurrence-based Genetic Screening), that enables large-scale GWAS on millions of *k*-mers in thousands of individuals for dozens of phenotypes. The relevance of the identified *k*-mers was evaluated by mapping them to gene-product annotations in public reference databases.

## Results

### Overview

We developed a *k*-mer-based GWAS approach that we used to screen for associations between gut-microbiome sequences and 26 host phenotypes (**Table S1**) in 938 healthy participants from the *Milieu Intérieur* study (**Fig. 1**). Gut-microbiome DNA was extracted and sequenced with whole-genome shotgun sequencing. From the shotgun data we built a *k*-mer abundance matrix of size *n* × *m*, where *m* is the number of distinct *k*-mers and *n* the number of individuals, using the kmtricks software^24^. Univariate GWAS of the *k*-mers were then performed with the MOGS software for every phenotype. In-silico functional analyses of the top-associated *k*-mers were carried out by systematic BLAST^25^ searches against the NCBI GenBank^26^ database to retrieve their potential functions. Finally, the results of the *k*-mer-based GWAS were compared with a standard taxa-based screening pipeline, in which taxa were profiled with Kraken2^3^ and MetaPhlAn4^4^, and functional annotations were estimated with HUMAnN^27^.

**Figure 1.**
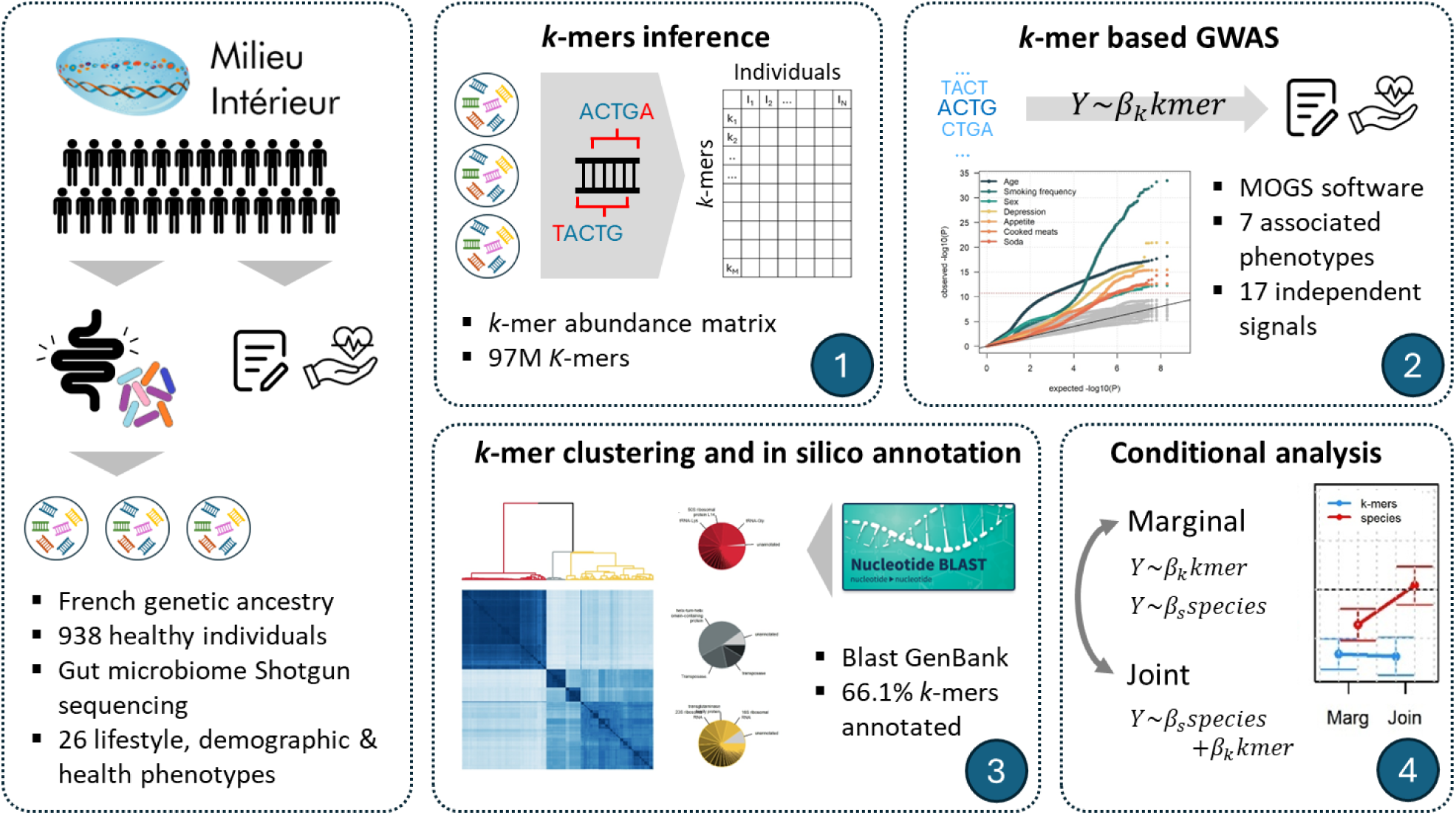
Study overview. We developed a *k*-mer based GWAS framework that we applied to 938 individuals from the Milieu Interieur cohort. Step 1 consists of the *k*-mer matrix computation and filtering using *kmtricks*. For step 2 we implemented MOGS, a software optimized for *k*-mer based GWAS, and executed it on 26 phenotypes. We provided on step 3 a clustering on the significant *k*-mers per phenotype and used *Blast* to annotate those clusters. In step 4, we conducted a comparison with taxonomy-based screening, comparing the relative contribution of *k*-mers and species using conditional analysis.

### Characteristics of k-mers derived from shotgun sequencing

We built a matrix of *k*-mer occurrences for the 938 participants of the *Milieu Intérieur* cohort using kmtricks^24^ with a *k*-mer length of *k* = 31, yielding 3,007,528,127 distinct *k*-mers. After filtering out k-mers that (i) had a maximum abundance below 20 across all samples of the dataset and (ii) were present in fewer than 5 % of individuals, 97,085,593 unique *k*-mers remained. The per-individual *k-*mer counts ranged from 140,766 to 34,012,926 (mean = 11,697,042; **Fig. 2a**) and were highly correlated with the number of reads per individual (**Fig. 2b**). This correlation motivated the use of relative *k-*mer abundances, i.e., normalizing each count by the individual’s total read depth. For comparison purposes, each *k-*mer was aligned to a bacterial taxonomic database at the species level using the *blastn* tool of BLAST^25^. In total, 64,143,091 *k-*mers (66.1 %) matched at least one bacterial species, covering 66,408 species that each had at least one hit. Among the *k-*mers that mapped to a species, 25 % (≈ 16 M) aligned to a single taxonomic identifier, whereas the remainder aligned to two (29 %), three (11 %), or ≥ four (35 %) species, yielding an average of 4.2 species per *k-*mer and a maximum of 448 species for a single *k-*mer (**Fig. 2c**). As expected, these highly multi-taxonomic *k-*mers largely correspond to conserved gene functions shared across many taxa (**Table S2**). We also examined the distribution of the number of unique *k-*mers per species (**Fig. 2d**) which was strongly proportional to its genome length (Pearson r = 0.77; **Fig. S1**). The observation that identical sequences are present in multiple species directly supports our central hypothesis: some of the microbiome genetic effects on the host may be captured more effectively with a *k-*mer-based approach than with traditional species-level analyses. Furthermore, the fact that 34% (N =32,942,502) of the *k*-mers cannot be mapped to any species highlights the potential of a sequence-based approach to capture signal from uncharacterized genetic components of the gut microbiome.

**Figure 2.**
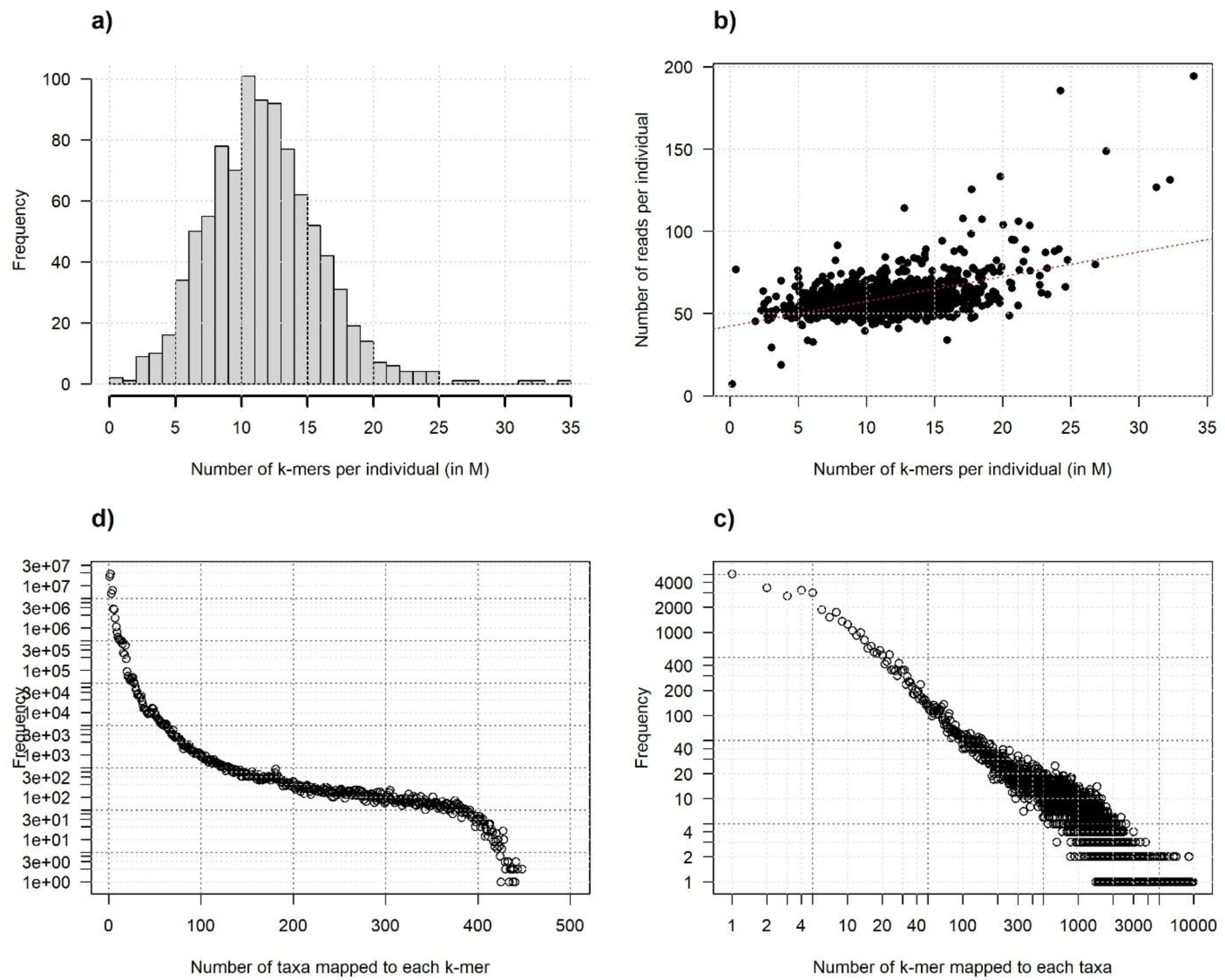
K-mer inference in the Milieu Interieur study. We computed a k-mer matrix for the 938 individuals from the Milieu Interieur cohort using *kmtricks*. Panel (a) presents the distribution of the number of distinct *k*-mers per individual. Panel (b) shows the number of distinct k-mers per individual as a function of the number of reads per individual. The red dash line indicates the regression coefficient (*ρ* = 0.5, *P* = 1.4 x 10^-70^). We mapped all *k*-mers to bacterial species in the NCBI database. A total of 64,143,091 *k*-mers map to at least one species. Panel c) and d) displays the distribution of the number of taxa mapped with each *k*-mer and the distribution of the number of *k*-mers mapped to each species in that subgroup.

We derived taxonomic profiles for the gut-microbiome sequencing reads of the 938 participants using Kraken2^3^ and MetaPhlAn4^4^. Kraken2 identified substantially more species than MetaPhlAn4^28^. Kraken2 inferred 1 074 unique taxonomic IDs (average ≈ 197 species per individual), whereas MetaPhlAn4 inferred 666 unique taxonomic IDs (average ≈ 91 species per individual). In total, 310 species were detected by both methods. Among these shared taxa the relative abundances showed strong agreement (mean Pearson r = 0.92 across all individuals; **Fig. 3a**). For each method we extracted the *k-*mers that map to the species identified by Kraken2 (1 074 taxa) and MetaPhlAn4 (666 taxa). The *k-*mers associated with Kraken-derived species accounted for 47 % of the total *k-*mer pool (N = 45 551 958), while those linked to MetaPhlAn-derived species represented 59 % (N = 57 076 670) (**Fig. 3b**). A small fraction of the species inferred by either classifier could not be linked to any of the extracted *k-*mers; however, when we restricted the analysis to the 310 overlapping species, the unmapped proportion fell to 1.6 % of all species (**Fig. 3c**). We then computed, for each species, the correlation between its relative abundance (as estimated by the taxonomic profiler) and the summed abundance of its associated *k-*mers (**Fig. S2**). The resulting correlations showed partial agreement, with mean Pearson r = 0.58 for MetaPhlAn4 and r = 0.70 for Kraken2. These results reinforce the added value of a *k-*mer-based screening: it can capture microbial signals that are missed by taxon-based approaches, either because the sequences cannot be assigned to a species or because the contribution of a species is fragmented across many individual *k-*mers.

**Figure 3.**
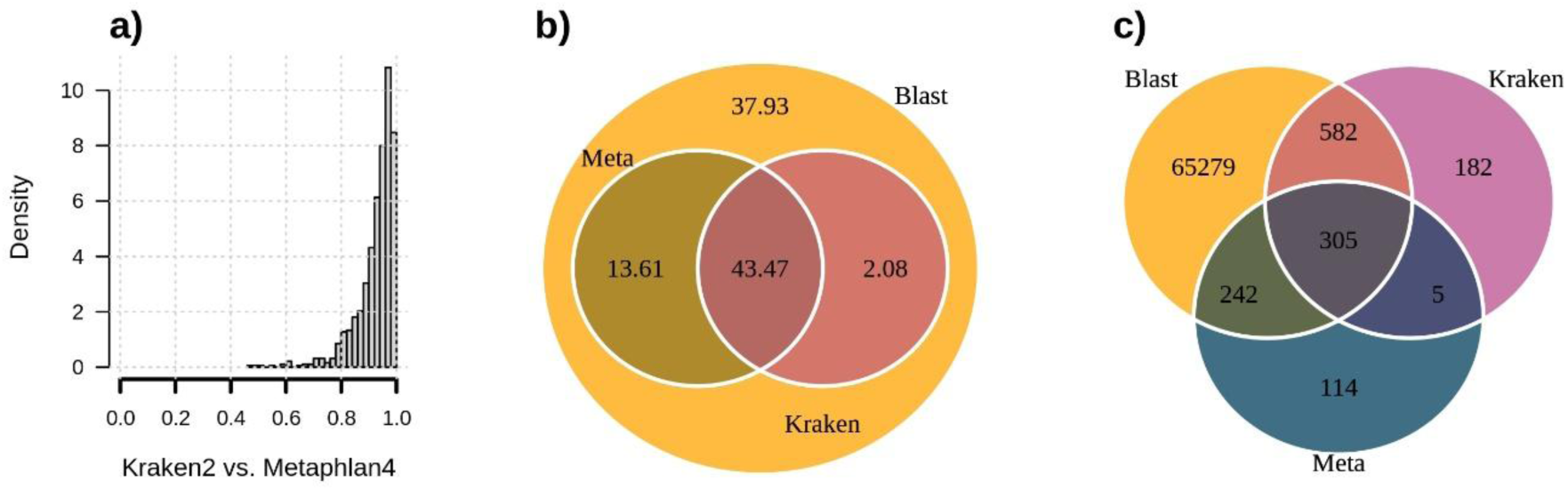
Comparison with Kraken2 and Metaphlan4. Panel (a) presents the distribution of the correlation of species’ relative abundances between Metaphlan4 and Kraken2 for a subset of 310 species quantified for both methods. Correlations were derived for each of the 938 individuals from Milieu Interieur as cor(Metaphlan4 abundance, Kraken2 abundance). Panel (b) shows a Venn diagram of the 97,085,593 *k*-mers inferred from *kmtricks*. Circles indicate whether the *k*-mers were part of the total pool of *k*-mers (*Blast*) mapped to species quantified in Metaphlan4 (*Meta*), or by Kraken2 (*Krak*). Numbers are provided in millions. Panel (c) shows a Venn diagram of the species identified by either a Blast on the *k*-mers, Metaphlan4 (*Meta*) or Kraken2.

### A k-mer based metagenome GWAS

We developed MOGS (Metagenome Occurrence-based Genetic Screening), a tool for association-screening of *k*-mers. MOGS fits a standard univariate linear regression between an outcome *Y* and a *k*-mer *K* while adjusting for a set of covariates *C*_*i*_ (*i* = 1 … *M*): *Y*∼*β*_*k*_*K* + ∑_*i*=1…*M*_ *β*_*i*_*C*_*i*_, where *β*_*k*_ is the effect of the tested *k*-mer and *β*_*i*_ is the effect of covariate *C*_*i*_. The MOGS implementation incorporates several algorithmic ingredients to optimize the runtime; a complete GWAS on all 97 M *k*-mers can be finished in < 25 min on a 20-core machine. To assess calibration under the null hypothesis, we used the 97,085,593 *k*-mers from the *Milieu Intérieur* cohort and simulated phenotypes that are independent of the microbiome. Two standard GWAS diagnostics were applied: (i) visual inspection of the quantile-quantile (QQ) plot of observed versus expected −log _10_ *P*values, and (ii) the genomic inflation factor (*λ*), which quantifies deviation of the median test statistic from its null expectation. As shown in **Figure S3a–b**, the QQ plots are well-behaved and the *λ* values are close to 1, indicating proper calibration. A slight deflation of the tail of the distribution was observed; this is likely explained by the extreme correlation among *k*-mers that arises from their overlapping construction (e.g., two 31-mers can share up to 30 nucleotides)^29–32^ (**Figure S3c**). To quantify the impact of this correlation we performed a clumping procedure across a range of *r*^2^ thresholds. Only 287 *k*-mers remained mutually uncorrelated at *r*^2^ ≤ 0.01 (**Figure S3d**; see **Supplementary Notes**), confirming that the effective number of independent tests is far lower than the number of k-mers (97 M).

We applied MOGS to the *Milieu Intérieur* cohort to perform genome-wide association studies (GWAS) on 26 phenotypes spanning demographics, physiological measurements, medical history, mental-health status, diet, physical activity and smoking habits (**Table S1**, **Fig. S4**). Using a strict Bonferroni-corrected significance threshold of 1.9 × 10⁻¹¹ (accounting for ≈ 2.5 billion tests), seven phenotypes showed significant associations with *k*-mers: age, sex, depression, smoking frequency, appetite, cooked-meat consumption, and soda intake (**Fig. 4a–b**). Previous studies have reported links between the gut microbiome and these traits at various levels (alpha- and beta-diversity, community composition, single taxa, etc.)^33–38^. Age exhibited the strongest overall enrichment for signal: 86 097 *k*-mers reached genome-wide significance, with a minimum p-value of 6.7 × 10⁻¹⁹. By contrast, the numbers of significant *k*-mers were lower for the other traits (smoking frequency = 5 310, depression = 1 965, appetite = 393). The lowest p-value among all phenotypes was observed for smoking frequency (3.4 × 10⁻³⁴). There was essentially no overlap in the sets of significant *k*-mers across phenotypes, except for a shared subset of 163 *k*-mers between depression and appetite. Nevertheless, the association statistics were highly correlated across the seven traits (**Fig. S5**). For instance, age displayed a strong negative genetic correlation with sex (r = -0.85, p < 10⁻³⁰⁰) and with depression (r = -0.59, p < 10⁻³⁰⁰), whereas sex and depression were positively correlated (r = 0.75, p < 10⁻³⁰⁰).

**Figure 4.**
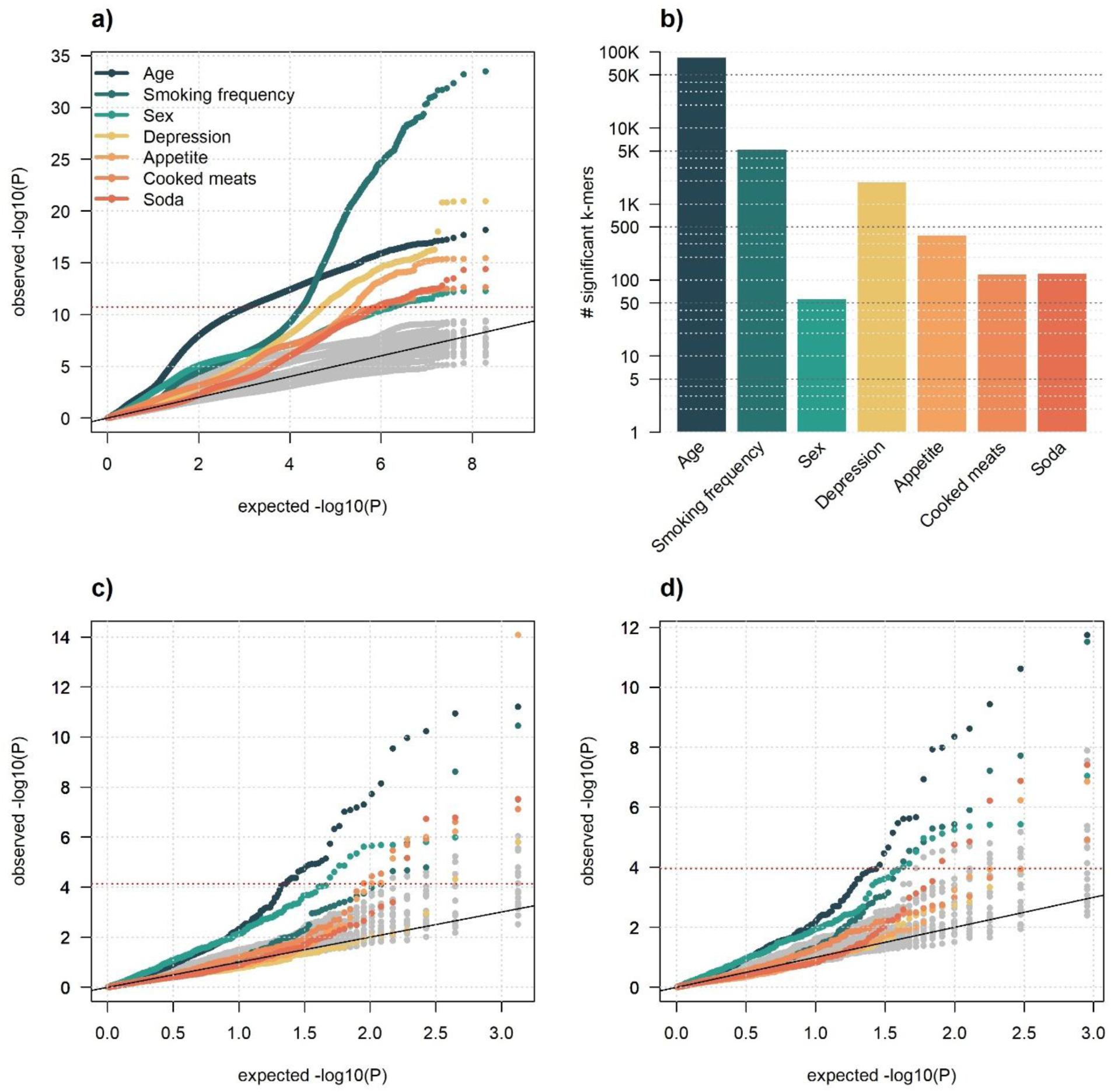
k-mer-based metagenome GWAS analysis in Milieu Interieur. Metagenome *k*-mer-based GWASs for the 26 phenotypes in the Milieu Interieur cohort. Panel a) presents the QQplot of all phenotypes highlighting the seven phenotypes with *k*-mers reaching a stringent Bonferroni corrected significance threshold (horizontal red dash line, P < 2 x 10^-11^). Panel b) shows the count of significant *k*-mers for those seven phenotypes. Panels c) and d) present the QQplot for the screening for association between the 26 phenotypes and the species relative abundance inferred by Kraken2 and Metaphlan4, respectively. The latter only includes species present in at least five individuals.

As illustrated in **Figures S6–S12**, the top-associated *k*-mers show a strong correlation structure, indicating that the thousands of significant hits actually represent only a limited number of independent-association signals (ISs). To estimate the number of ISs we performed a stepwise conditional-regression analysis (see **Methods**). Briefly, for each of the seven associated phenotypes we built a joint model that started with the most significant *k*-mer and then added any further *k*-mers that remained significant after conditioning on those already in the model. Overall, we identified 17 ISs. Appetite and depression each harbored four ISs; Smoking frequency and cooked-meat consumption each harbored three ISs; and Age, sex, and soda consumption each harbored a single IS (**Table S3**). The top *k*-mers were generally rare (present in < 15 % of individuals), except for the age-associated *k*-mers (present in > 99 % of samples) and the sex-associated *k*-mers (present in > 91 %). They mapped to a relatively small set of species (average *N̄* = 7.1 species per *k*-mer), with the notable exception of the age-associated *k*-mers, which mapped to 107 species—consistent with these *k*-mers originating from core-genome regions shared across many taxa.

### In-silico functional analysis highlight microbiome functions

To explore the possible functional roles of the phenotype-associated *k*-mers, we performed a series of in-silico functional annotations by mapping the *k*-mers with potential protein functions to the NCBI GenBank database^26^. On average, 79 % of the associated *k*-mers could be assigned to a bacterial species, and 64 % could be linked to a gene product within that species (**Table S4**). The species and functional annotations were highly overlapping within each phenotype, resulting in a limited number of distinct entities (average *N̄*_taxID_ = 114 taxa and average *N̄*_function_ = 24 functions per phenotype), except for age, which mapped to > 1 919 taxa and > 440 functions. To account for the strong correlation among *k*-mers and to test for enrichment of specific annotations, we clustered *k*-mers on the basis of their pair-wise correlation and then summarized the functional annotations for each cluster (**Table S5**). **Figures S6–S12** show, for each phenotype, the significant *k*-mers, clustered based on their correlation, the dominant annotations within each cluster, and the location of the independent association signals (ISs), indicating that for several clusters, the observed associations tend to correspond to biologically distinct functions. For easier downstream follow-up, we also extracted, from each cluster, the *k*-mer whose functional annotation best represents the cluster’s overall annotation (**Table 1**, **Table S6**).

**Table 1.**
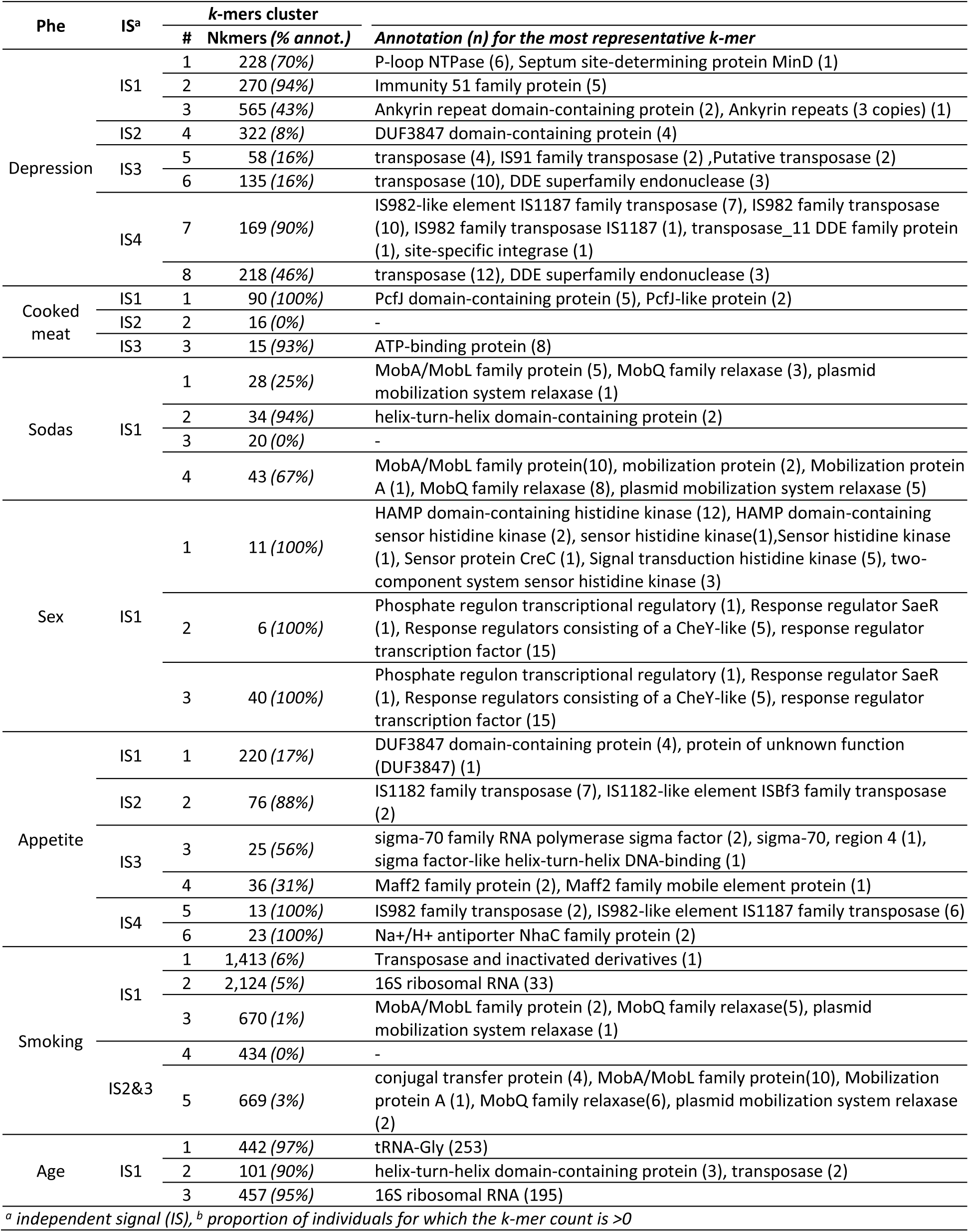
Top independent associated k-mers.

The enrichment of functional annotations in several *k*-mer clusters was homogeneous, suggesting that a particular biological mechanism may underlie the association. The most intriguing example was cluster 3 of the depression phenotype, whose *k*-mers map almost exclusively to genes containing ankyrin-repeat domains (ANKRD) (**Fig. S8**). Ankyrin-repeat motifs are common structural elements in proteins from both bacteria and eukaryotes, but they are far more abundant in eukaryotic genomes. In humans, several ANKRD genes—TRANK1, ANK1/2/3, and SHANK1/2/3—are well-established risk factors for a range of neuro-psychiatric disorders^39–41^. Prior work has shown that *host* mutations in ANKRD genes can influence gut-microbiome composition^42^, and conversely, that fecal-microbiota transplantation experiments in mice can induce depression-like behaviours, possibly through modulation of host TRANK1, SHANK1, and SHANK3 expression^43,44^. Our results extend these observations by suggesting that bacterial ankyrin-repeat-containing genes present in the gut microbiota may also affect mental-health phenotypes in the host. Importantly, host-DNA contamination was removed prior to all analyses (see **Methods**). As an additional control, we also BLASTed the 240 *k*-mers annotated as ANKRD; none of them matched the *Homo sapiens* NCBI database.

A few additional annotations merit further investigation. Sex-associated *k*-mers clustered into three groups, all of which were annotated (**Fig. S11**). The overwhelming majority of these annotations were related to the two-component system (TCS) histidine-kinase pathway, with the two most frequent terms being “histidine kinase” and “response regulator”. The annotation “response regulator, CheY-type” further points toward chemotaxis and motility via the CheA-CheY phosphotransfer complex^45^. Differences in gut-microbiota composition between males and females are well documented^46^; sex hormones have been implicated in those differences^47^, and experimental work in pigs has linked the TCS pathway and chemotaxis to host androgen levels^48^. Cluster 6 from the appetite-associated *k*-mers (**Fig. S7**) also showed a striking enrichment for “Na⁺/H⁺ antiporter NhaC family protein” (96.2 %). NhaC antiporters are membrane proteins that regulate intracellular pH and Na⁺ homeostasis. They have been extensively studied in *Bacillus* species^49,50^, including the probiotic *Bacillus subtilis*^51^, which has previously been associated with appetite regulation in animal models^52,53^. Finally, age-associated *k*-mers mapped to a very large and heterogeneous set of annotations (**Fig. S6**). Two broad functional categories were disproportionately represented: (i) ribosomal proteins (50 S, 30 S, 23 S, SSU, etc.; 41.3 %) and (ii) transfer RNAs (tRNA-Gly, tRNA-Lys, tRNA-Met, etc.; 36.5 %). Although this pattern may reflect age-related shifts in bacterial metabolic activity, it could also be a confounding effect: these housekeeping genes may act as proxies for overall gut-microbiota composition rather than causal drivers.

### Contribution of MOGS as compared to species-based screening

For comparison, we performed a conventional species-based GWAS on the same 26 phenotypes using the taxonomic profiles generated by Kraken 2 and MetaPhlAn 4. Only species that were detected in at least five individuals were retained (**Tables S7-S8**). The quantile-quantile (QQ) plots from both taxonomic-based screens display the same pattern as the *k*-mer-based GWAS, with enrichment of associations for the same traits—age, sex, smoking, cooked-meat consumption, and soda intake (**Fig. 4c–d**). After Bonferroni correction, 31 Kraken2 species-phenotype pairs and 24 MetaPhlAn4 species-phenotype pairs passed the significance threshold (**Tables S7-S8**). Apart from smoking status, which was linked to two species, no additional phenotypes were uncovered by the species-based approach, indicating that sequence-based and species-based screenings have comparable power to detect phenotype-associated signals.

We next examined, for each phenotype, how the 17 IS *k*-mers correlate with individual taxa inferred by Kraken2 (**Fig. 5a**) and MetaPhlAn4 (**Fig. 5b**). As expected, most species show only modest correlations with these *k*-mers (average absolute ρ = 0.031 for Kraken 2-derived taxa and 0.028 for MetaPhlAn 4-derived taxa). Nonetheless, several species display both a substantial correlation (ρ > 0.5) and a statistically significant association with the same phenotype (**Table S9**). The strongest concordance was observed for the IS2 *k*-mers linked to *smoking frequency*: they correlate with *Longicatena caecimuris* (ρ = 0.74) and this bacterium is itself associated with smoking frequency (Kraken2 p = 3.5 × 10⁻¹¹; MetaPhlAn4 p = 3.0 × 10⁻¹²). A few other IS *k*-mers also show ρ > 0.5 with a species that is significantly associated with the same outcome (e.g., IS1 for age and IS1 for appetite), indicating overlapping signals. To determine which entity is the primary driver of the association, we performed conditional analyses. For each phenotype we compared the regression coefficient from a marginal model (including only the top IS k-mer or only the associated species) with that from a joint model that contains both variables. As illustrated in **Figure 5c–d**, adding the associated *k*-mer to the model containing the species causes a pronounced attenuation of the species’ effect size, driving most species’ associations to the null (all but one become non-nominally significant). This demonstrates that, in our data set, the top associated *k*-mers can explain a large fraction—and in some cases the entirety—of the species-phenotype association.

**Figure 5.**
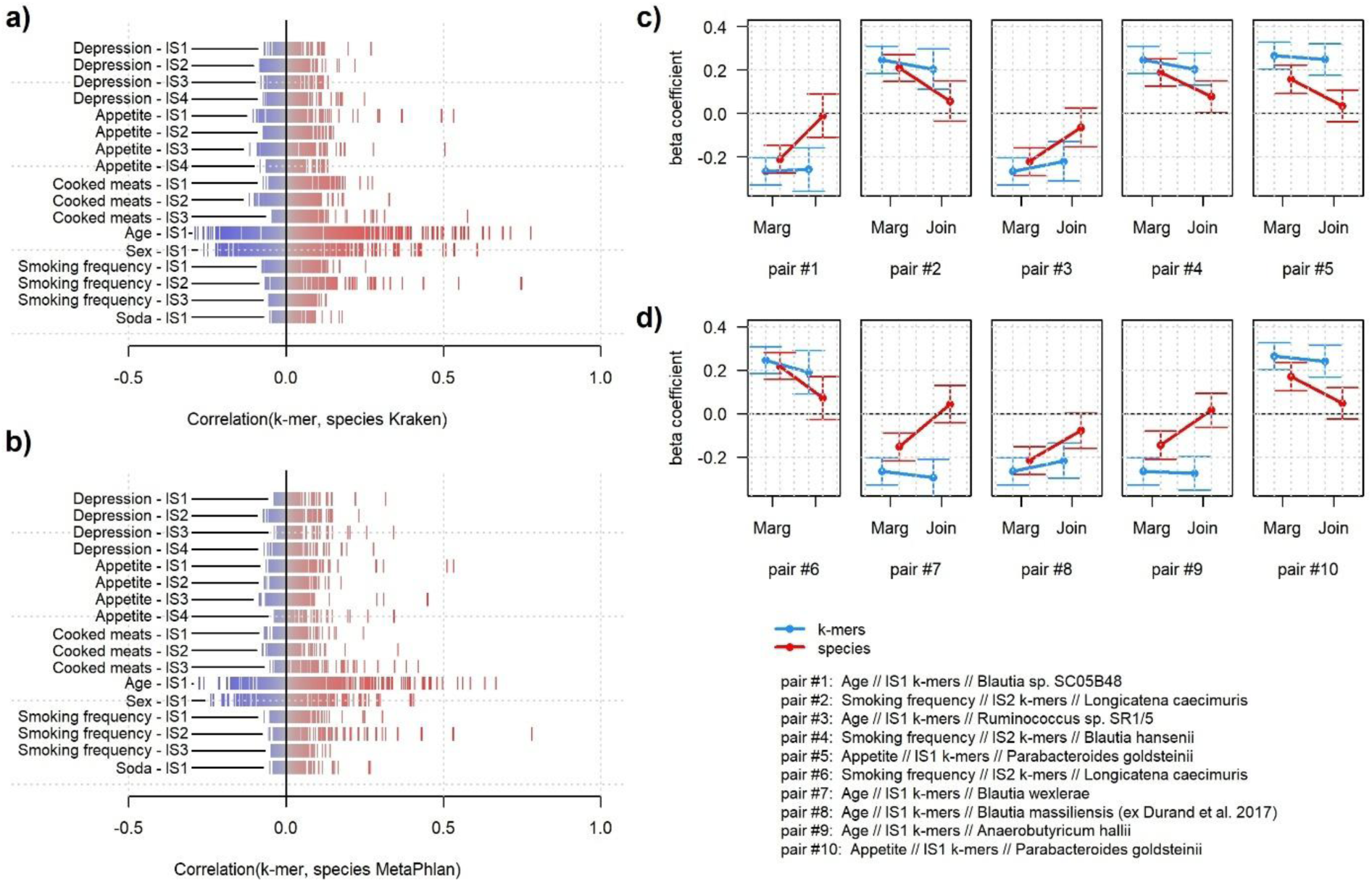
Correlation between top k-mers and bacteria. Correlation between the 17 IS *k*-mers (**Table S3**) and bacteria species estimated from MetaPhlan4 (a) and Kraken2 (b). Each row represents a single IS *k*-mer and its associated phenotype. For the top five IS *k*-mers -species pairs showing both high correlation and significant association with the same phenotype, we compared estimates from marginal model (Y ∼ *k*-mers and Y ∼ species) against estimates from a joint model including both variable (Y ∼ *k*-mers + species). Panel (c) presents those results for the top pairs from Kraken2, and panel (d) for the top pairs from MetaPhlan4. For instance, pair #1 compares the estimates between the IS1-age *k*-mer and the *Blautia species SC05B48*, which displays the largest correlation with the IS1-age *k*-mer (ρ = 0.78) and significantly associated with age (P = 5.8 x 10^-11^).

Finally, we conducted an *in silico* functional enrichment based on the species using HUMAnN^27^ and MaAsLin3^54^ on all 26 phenotypes. Briefly, HUMAnN built a relative abundance matrix of genes from the sequencing metagenome reads for each individual from the cohort, and the abundance is then tested for association with the phenotype using the regression model from MaAsLin3. The gene abundance matrix included 118,628 genes and the MaAsLin3 regression pointed to thousands of potential functions and pathways when using the default false discovery rate threshold (**Table S10**), making it challenging to draw any conclusion about the mechanisms involved. However, when using a more stringent Bonferroni correction threshold to focus on key annotations, only age and sex revealed significant enrichment, with respectively 14 and 10 associated genes (minimum *P*-values of 5.53 x 10^-14^ and 5.10 x 10^-10^, respectively) (**Table S11**, **Fig. S13b**). As expected, genes under the false discovery rate threshold are considerably more important than average (**Fig. S13a**). There was almost no overlap between those functions and those identified in the *k*-mers based analysis, suggesting that the two approaches may be capturing different signals.

## Conclusion

The gut microbiota contribute a wide range of functions to the host, including nutrient provision, protection against infection, and immune development^55^. Although individual isolates can harbor uniquely specific functions, most reported associations between human phenotypes and taxonomic groups point to functional classes that are shared by many species or to broader community-level characteristics such as taxonomic diversity^56^.

In this work we build upon the simple hypothesis that some of the links between human traits and the gut microbiome are driven by gene functions that are shared across multiple taxa. To test this hypothesis, we built a taxonomy-free metagenome-wide association study (GWAS) that screens for associations between *k*-mers derived from shotgun sequencing reads and phenotypes of interest. By design, the approach captures variation not only in the core genome but also in gene-presence/absence and the entire accessory genome. Applying the method to 938 healthy participants of the *Milieu Intérieur* cohort, we first confirmed that a large majority of microbial DNA sequences are shared across several species. Screening 97M *k*-mers for association with 26 phenotypes revealed significant associations for seven traits. In cases where a species-based signal overlapped a *k*-mer-based signal, conditional analyses showed that the *k*-mer better explained the association, further suggesting that shared sequences may indeed better specify the mechanism involved.

Sequence-centric analysis are not without limitations. Here, we focused on two main challenges. First, the interpretation of *k*-mers hits is non-trivial, especially while functional databases are still incomplete. Our in-silico functional annotation identified plausible gene-product candidates for a few of the significant *k*-mers. The most intriguing result was the enrichment of ankyrin-repeat-domain (ANKRD) genes from the microbiome among *k*-mers associated with host depression. One possible mechanism is the translocation of ankyrin-repeat-containing proteins into host cells^57^. However, this particular enrichment, as well as other candidate mechanisms we report, are hypothesis-generating at most. They will need replication in independent cohorts and experimental validation. A second limitation is the extreme correlation among *k*-mers. For example, the age-related signal spreads across ∼80 000 highly correlated *k*-mers, making it difficult to pinpoint the causal sequence. A natural extension of the current workflow to address this question would be to assemble correlated *k*-mers into more contiguous sequences, thereby reducing redundancy and improving interpretability. Finally, another perspective is the exploration of the microbiome “dark matter” –*k*-mers that cannot be mapped to any known bacteria but were found associated with the phenotype study.

Overall, our work not only offers a proof of concept for the feasibility of sequence-based gut metagenome GWAS, but also opens new avenues for follow-up analyses that leverage methodologies developed for human GWAS. This includes quantifying the contribution of the microbiome to the total phenotypic variance using variance-component models developed in human genetics for heritability estimation, conducting screening for microbiota gene–gene and gene–environment interactions, and developing *k*-mers risk score of host phenotype using penalized models.

## Methods

### Milieu Intérieur: a gut microbiome cohort of 1000 individuals

The Milieu Intérieur consortium is a French cohort of healthy individuals stratified equally by decade, with age ranging from 20 to 69 years old. The consortium was established in 2012 with the objective of characterizing and understanding the genetic and environmental variabilities of human phenotypes based on gut microbiome data. A broad range of variables were collected, including genetic, genomic, and environmental data, on most participants. On their first visit the volunteers were also asked to fill in an extended form about socio-demographic, lifestyle and family health history, all recorded in an electronic case report form (eCRF). Gut microbiota from all volunteers were sequenced using shotgun metagenomic sequencing at Biotrial Inc. in Rennes, France. Reads mapping to human genome were removed. (see **Supplementary Notes**). A complete description of the cohort is available in Thomas et al 2015^58^. In this analysis we used 938 individuals, including 495 men and 443 women, with complete microbiome and phenotypic data.

### K-mers inference

Raw shotgun sequencing in BZIP2 format were transformed in gzip file and were pre-processed using Fastp^59^ to remove adapters and filter out reads that did not match standard quality control checks. The derivation of the *k*-mer abundance matrix from the clean gzip file and the filtering of low abundant *k*-mer was performed using *Kmtricks*^24^ and *k*=31, a commonly used length for *k*-mers analyses^60^. *Kmtricks* works in two steps. In the first stage, *Kmtricks* enumerates, deduplicates and counts all *k*-mers present in the input files. To decrease computational time this computation is done across multiple threads resulting in *k*-mer enumeration split over multiple partitions. During that stage, *Kmtricks* can apply various filtering on the *k*-mers. Here we discard *k*-mers with abundance below 20 in all individuals allowing to both speed-up the total running time and remove most of the sequencing errors. In the second stage, *Kmtricks* aggregates all partition files into a single complete matrix file. Overall, we obtain a total of 3,007,528,127 *k*-mers for a total disk space of 4.8To. We further filtered all *k*-mers shared by less than 5% of the individuals (N < 47) using a shell awk command resulting in a total of 97,085,593 *k*-mers, that is 3.22% of the original matrix. The final *k*-mer abundance matrix uses 184 GB of disk space. In terms of computational resources, we used 50 threads and 16.2 GB peak memory, for a wall-clock running time of about 6h30min at stage 1, and 50 threads and 371 MB peak memory for a wall-clock running time of about 3d20h30min at stage 2.

### k-mers annotation

We mapped each and every *k*-mers to bacteria with the *blastn* tool of BLAST^25^ on a local instance of the nucleotide database (version 1.2). Default settings were applied. Although BLAST requests are fast, they have high memory footprint. We sub-divided our 97 million sequences into smaller batches. After benchmarking, the batch size was set to 1 million sequences per run. It leads to a 250 GB peak memory per run. To improve running-time, the requests were parallelized over 30 CPUs and used only the bacterial genomes NCBI reference database (*-taxids 2* option).

In order to run an exhaustive functional screening of thousands of *k*-mers, we implemented an API of *Blastn,* given the accession numbers and the starts and ends of alignments in our locally computed BLAST output, to access the GenBank of NCBI. Because of the computational constraint, we investigated the functional properties of the significant *k*-mers from the seven phenotypes only. Due to the strong enrichment of the 86,097 significant *k*-mers of age with species in the BLAST database (1,147,738 associations), we further restrained the analysis to the top 1,000 *k*-mers, resulting in 45,269 associations. The functional analysis gave 40,910 gene/coding proteins for age (848 *k*-mers were mapped to a gene), 8429 for depression (1,321 *k*-mers), 6406 for smoking habits (127 *k*-mers), 1,231 for sex (57 *k*-mers), 1,001 for appetite (160 *k*-mers), 419 for cooked meat consumption (112 *k*-mers) and 308 for sodas (76 *k*-mers) (**Table S4-5**).

### k-mer based Metagenome GWAS

We conducted metagenome genome-wide association studies (GWAS) on a set of 26 phenotypes stratified in 7 categories: demographics, basic physiological measurements, psychological problems (sleep, drug, …), smoking habits, medical history, food and nutrition, and socio-professional information. Further details on the phenotypes are available in **Table S1**. We adjusted each GWAS for age and sex. For age and sex phenotypes, we corrected respectively for sex and BMI, and age and BMI. The association between each *k*-mer and the outcomes considered was assessed using a standard linear model. Let *n*, *m* be respectively the number of individuals and *k*-mer. Let ***y*** be a vector of a categorical or a quantitative phenotype, ***X*** be a *n* × *m* matrix of *k*-mer abundances, and ***C*** a *n* × 𝑣 matrix of covariates factors, where 𝑣 represents the number of covariates. We corrected each row of ***X*** by the number of short reads of its associated individual and centered-reduced ***y***, ***X***, ***C***. Our linear model is then of the form:

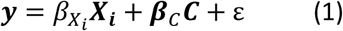

with *β*_*X*_*i*__ the effect of the i^th^ *k*-mer ***X***_***i***_, ***β***_*C*_ the vector of effects of the covariates and ε the standard error. The significance of *β̂*_*X*_*i*__, the estimated *k*-mer effect, is derived using Wald test. We applied a stringent Bonferroni correction for multiple testing, corresponding to a threshold of *p* < 1.91 x 10^-11^ (0.05/(27 × 97,085,593)) to assess *k*-mer significance.

### Software implementation

To address the computational challenges of the metagenome GWAS, we used the following algorithm speed-up. For simplicity, we assume that all variables are standardized to have mean 0 and variance 1. First, all marginal correlations are derived:

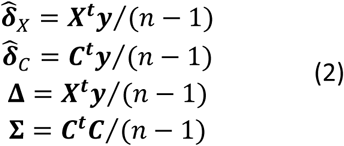

where *δ*_*X*_ is an 1 × *m* vector of *k*-mer correlation with ***y***, *δ*_*C*_ is an 1 × 𝑣 vector of correlation between the covariates and ***y***, ***Δ*** is an *m* × 𝑣 matrix of *k*-mer correlation with the covariate, and ***∑*** is an 𝑣 × 𝑣 matrix of covariate correlation. Second, for each *k*-mer *i*, we build from the marginal correlation (eq. 2), the covariance matrix of the joint effects model (eq. 1):

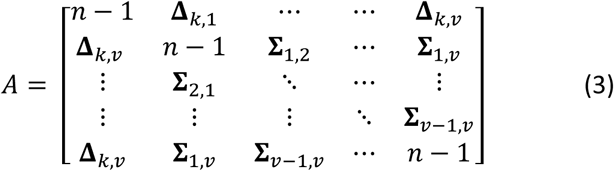

Third, we derive for each *k*-mer the coefficients from the joint model (eq. 1) from the marginal coefficient *δ̂* = (*δ̂*_*X*_*i*__, *δ̂*_*C*_) and the inverse of the covariance matrix:

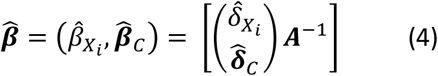

Finally, the *P*-value for significance of *β̂*_*X*_*i*__ is estimated from a 1 degree-of-freedom chi-squared distribution at 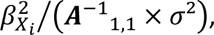 where 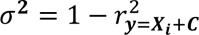 is the residual variance of ***y*** from the joint model (eq. 1) derived as 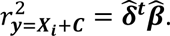 The above formulation was implemented in C++ in the MOGS package.

To keep running memory low, MOGS processes the input *k*-mer matrix in chunks of 100,000 *k*-mers, and each chunk is read and processed in parallel to minimize runtime. This keeps the RAM usage to less than 3 Gigabytes for the whole analysis. The software has approximately linear complexity regarding the size of the input matrix (**Fig. S14**).

### Enumeration of independent k-mers

For each phenotype associated with multiple *k*-mers, we implemented a stepwise selection procedure to estimate the number of independent association and build a joint association model defined as: ***y*** ∼ ∑_*i*∈Ω_ *β*_*X*_*i*__***X***_*i*_ + ***β***_*C*_***C***, where ***C*** is a matrix of covariates and Ω is a set of *k*-mers significant below the Bonferroni significance threshold (*p* < 1.91 x 10^-11^) in this joint model. The procedure worked as follows: i) we set a baseline model by adding the most associated *k*-mer ***X***_*i*_ to the set Ω = *i*. ii) all remaining *k*-mers ***X***_*i*∉Ω_ are tested for association one-by-one in a joint model: ***y*** ∼ ∑_*i*∈Ω_ *β*_*X*_*i*__***X***_*i*_ + *β*_*X*_*i*∉Ω__***X***_*i*∉Ω_ + ***β***_*C*_***C***, and the association *p*-value *p*_*i*∉Ω_ for each ***X***_*i*∉Ω_ is derived iii) if no *p*_*i*∉Ω_ is below the Bonferroni significance threshold (*P* < 1.91 x 10^-11^), the procedure stops. Otherwise, the most significant *k*-mer (i.e. *p*_*i*∉Ω_ = min(*p*_*i*∉Ω_)) is added to Ω. iv) step ii) to iv) are repeated until there are no additional *k*-mer can be added to Ω.

### Inferring Species from shotgun data

We inferred species abundance using two taxonomic classifiers, Kraken2^3^ and MetaPhlAn4^4^, focusing on the species level. Kraken2 generates all the *k*-mers from the input sequences and aligns them to a predefined taxa database. It has heavy memory usage because it loads the entire reference taxonomic database during the inference process, leading to memory issue (requiring over 1To of RAM when using the ‘nt’ database). We therefore reduced the NCBI reference database ‘nt’ to a local downloaded database ‘kraken_standard’ which focuses on metagenomic bacteria species of the NBCI database. To improve the running-time, we further parallelized the inference over 5 CPUs. We remove the unclassified reads (∼45.63%), the uncultured species (∼0.5%) and the species whose relative abundance is less than 0.01% (∼97.5%) to reduce the false positive rate issue. MetaPhlAn4 is a solution to avoid some of the Kraken2 issues. It aligns the sequences to a predefined database of marker genes^61^. This method is more conservative than Kraken2, and keeps only the core of the taxonomic profile^62^. However, it is computationally slower and is less relevant for short sequences. To improve the running-time, we parallelized over 10 CPUs. We remove the unclassified species (∼26.9%) and the species whose taxonomic ID is not reported (∼28.4%).

### Species-based screening

We conducted univariate species association screening on a set of same 26 phenotypes used in the *k*-mer GWAS. When needed, we removed individuals without the information of a given phenotype. All species were tested using a standard univariate linear regression, defined as: ***y*** = *β*_*S*_*i*__ ***S***_***i***_ + ***β***_*C*_***C*** + ε, where ***y*** is a vector of a categorical or a quantitative phenotype, ***S*** is a *n* × *p* matrix of species relative abundance, and ***C*** is a *n* × 𝑣 matrix of covariates factors, where 𝑣 represents the number of covariates. We adjusted all models for age and sex. For age and sex phenotypes, we corrected respectively for sex and age. As for the *k*-mers, we applied a stringent correction for multiple testing accounting for the number of phenotypes and the number of taxa tested, resulting the P-value thresholds of Pt_kraken_ = 1.8 x 10^-6^ and Pt_metaphlan_ =2.9 x 10^-6^.

### Species annotation

We annotated the taxonomic profile of each individual using HUMAnN 3.0^27^ from our raw sequencing data, and applied MaAsLin3^54^ to the resulting annotation to assess the enrichment for specific annotation conditional on individuals’ phenotypic values. Briefly, HUMAnN constructs the taxonomic profiles within a dataset from the metagenome sequencing reads, and determines an abundance of each gene family in the community of bacteria species for each single individual. Here we built the annotation based on the UniProt Reference Clusters (UniRef50) database. MaAsLin3 was then used to perform a standard univariate linear regression between each specific protein annotation ***G***_g_ and a phenotype ***y*** using the following model: ***G***_g_ = *β*_y_***y*** + ***β***_C_***C*** + ε, where *β*_y_ is the regression coefficient for phenotype ***y***, ***C*** is a matrix of covariates, and ***β***_C_ is the associated vector of covariate’s effect. We applied MaAsLin3 for the 26 phenotypes and adjusted all analyses for age and sex, except when age or sex were used as the predictor of interest.

### k-mer and species joint modelling

There were multiple instances where an independent signal (IS) *k*-mer and a bacteria species display correlation larger than 0.5 and were significantly associated with the same phenotypes, suggesting the two associations were capturing the same signal. To assess the relative fitness of either variables (*k*-mer or species) in explaining the phenotype, we conducted conditional analysis. For top 5 instances from Kraken2 and MetaPhlan4 species quantification, we compared the regression coefficient from a marginal model including only the top IS k-mer (***y*** = *β*_*X*_*i*__***X***_***i***_ + ***β***_*C*_***C***) or only the associated species (***y*** = *β*_*S*_*i*__ ***S***_***i***_ + ***β***_*C*_***C***) with that from a joint model that contains both variables (***y*** = *δ*_*X*_*i*__***X***_***i***_ + *δ*_*S*_*i*__***S***_***i***_ + ***β***_*C*_***C***). **Figure 5c-d** display these comparisons, showing *β*_*X*_*i*__ and *β*_*S*_*i*__ against *δ*_*X*_*i*__ and *δ*_*S*_*i*__ along their 95% confidence interval.

## Data availability

The dataset used to compute the *k*-mer matrix and the individual metadata are available in the European Genome-Phenome Archive under accession code EGAS00001004437.

## Code availability

The MOGS software is freely available at https://gitlab.pasteur.fr/statistical-genetics/MOGS. We used other tools that were previously published, and are listed below for convenience. The *k*-*mer*-matrix construction is done with the *kmtricks* tool (https://github.com/tlemane/kmtricks). The linkage between *k*-mer sequences and bacteria genomes/families is based on Blast version 2.16.0 (https://github.com/blast-io/blast). Gut microbiome taxonomy was done using MetaPhlAn4 and Kraken2 (https://github.com/biobakery/MetaPhlAn and https://github.com/DerrickWood/kraken2).

## Acknowledgements

This research was supported by the Agence Nationale pour la Recherche (ANR-20-CE15-0012-01) and the INCEPTION program (Investissement d’Avenir grant ANR-16-CONV-0005). The Milieu Interieur consortium was also supported by the Agence Nationale pour la Recherche (ANR-10-LABX-69-01). The funders had no role in study design, data collection and analysis, decision to publish, or preparation of the manuscript.

## Author information

These authors jointly supervised this work: Rayan Chikhi, Hugues Aschard.

## Authors and affiliations

**Institut Pasteur, Université Paris Cité, Department of Computational Biology, F-75015 Paris, France**

Raphaël Malak, Arthur Frouin, Léo Henches, Antoine Auvergne, Rayan Chikhi & Hugues Aschard

**Sorbonne Université, INSERM, Centre de recherche Saint-Antoine, CRSA, Microbiota, Gut and Inflammation Laboratory, Hôpital Saint-Antoine (UMR S938) Sorbonne Université, 27 rue Chaligny, 75012 Paris, France**

Harry Sokol

**Paris Center for Microbiome Medicine, Fédération Hospitalo-Universitaire, 184 rue du Faubourg Saint-Antoine, 75571 PARIS Cedex 12, France**

Harry Sokol

**Gastroenterology Department, AP-HP, Saint Antoine Hospital, 184 rue du faubourg Saint-Antoine, 75012 Paris, France**

Harry Sokol

**INRAE Micalis & AgroParisTech, UMR1319, Micalis & AgroParisTech, 4 avenue Jean Jaurès, 78352 Jouy en Josas, France**

Harry Sokol

**Program in Genetic Epidemiology and Statistical Genetics, Harvard T.H. Chan School of Public Health, Boston, MA, 02115, USA**

Hugues Aschard

## Contributions

R.C. and H.A. conceived the study and A.F., R.C., H.A. and R.M. designed research. H.A. acquired funding for the project. H.A. and R.M. wrote the original manuscript with contribution from A.F., L.H., H.S. and R.C. *Milieu Intérieur Consortium* provides the data. R.C. and R.M. contributed to sample collection and *k*-mer matrix generation. L.H. and R.M. implemented MOGS software. A.F., A.A. and C.B. contributed addressing methodological questions. R.M. performed the *k*-mer-based GWAS with support from A.A., C.B. and H.A. H.A. and R.M. performed Kraken2 and MetaPhlAn4 analysis and comparison. R.M. implemented the BLAST API with support from L.H., and performed HUMAnN and Maaslin3 analysis with support from H.S and H.A.- L.H. H.A. and R.M. contributed to the clustering, the independent signal analysis and the clumping procedure to account for *k*-mer correlation. All authors reviewed and approved the manuscript.

## Corresponding authors

Correspondence and requests for materials should be addressed to Raphaël Malak, Rayan Chikhi or Hugues Aschard.

## Consortia

### Milieu Intérieur Consortium

Laurent Abel, Andres Alcover, Hugues Aschard, Philippe Bousso, Nollaig Bourke, Petter Brodin, Pierre Bruhns, Nadine Cerf-Bensussan, Ana Cumano, Christophe D’Enfert, Ludovic Deriano, Marie-Agnès Dillies, James Di Santo, Gérard Eberl, Jost Enninga, Jacques Fellay, Ivo Gomperts-Boneca, Milena Hasan, Gunilla Karlsson Hedestam, Serge Hercberg, Molly A. Ingersoll, Olivier Lantz, Rose Anne Kenny, Mickaël Ménager, Frédérique Michel, Hugo Mouquet, Cliona O’Farrelly, Etienne Patin, Sandra Pellegrini, Antonio Rausell, Frédéric Rieux-Laucat, Lars Rogge, Magnus Fontes, Anavaj Sakuntabhai, Olivier Schwartz, Benno Schwikowski, Spencer Shorte, Frédéric Tangy, Antoine Toubert, Mathilde Touvier, Marie-Noëlle Ungeheuer, Christophe Zimmer, Matthew L. Albert, Darragh Duffy & Lluis Quintana-Murci

## Ethics declarations

### Competing interests

The authors declare no competing interests.

## Notes

### Competing Interest Statement

The authors have declared no competing interest.

