## Supplemental Figures for "Taxonomic-free metagenome GWAS to identify gut microbiome functions influencing host phenotypes"

**Supplementary Material**

### Supplementary Notes

#### Milieu Interieur gut microbiome data

Stool samples were collected during two visits. For participants that had two samples, we only kept the sample from the first visit. Shotgun metagenomic was applied to derive the bacterial profiles up to the species level. Detailed description of the data generation can be found in Byrd et al^1^. In brief, stool specimens were collected in a double-lined sealable bag containing a GENbag Anaer atmosphere generator (Aerocult; Biomerieux). Then fresh samples were aliquoted into cryotubes and stored at −80°C. Stool aliquots were shipped to the CRO Diversigen for DNA extraction and shotgun metagenomic sequencing using an Illumina HiSeq 2500. In the end, 21 trillion raw paired-end reads from 1,359 samples from 946 of the donors were obtained. To process the reads, Illumina TruSeq adapters were trimmed with Trimmomatic v0.36^2^; low-quality and low-complexity reads were removed with prinseq-lite 0.20.4^3^; and Bowtie2 v2.1.0^4^ was used to remove reads mapping to PhiX or the PacBio human genome. After processing, there were on average 13.9 ± 2.9 million reads per sample. Of an initial 1,000 recruited donors, 44 were excluded from this analysis because of lack of consent for sharing their data outside of the MI consortium. An additional 10 donors were excluded because of technical issues in the extraction and sequencing steps (e.g., low DNA extraction yield), resulting in a sample size for the shotgun dataset of 946 donors. In this analysis, we further removed eight individuals whose ID were not well registered, leading to a cohort of 938 individuals.

#### Correlated *k*-mers analysis and clumping procedure

To highlight the effect of high correlation between *k*-mers on the QQplot, we implemented a selection procedure to extract a group of *k*-mers that display particularly large correlation ($r^{2}$ $>0.5$). The selection procedure worked as follows: i) we randomly selected 1,000 *k*-mers, referred further as anchor *k*-mers across the entire set of 97,085,593 *k*-mers. ii) For each of those 1,000 *k*-mers, we derived their squared-correlation with all other *k*-mers and kept only those who display an $r^{2}$ $>0.5$ with at least one anchor *k*-mers. iii) This resulted in a set of 151,520 *k*-mers for which we extracted the *p*-value from each of the 50 null model simulations. The distributions of the *P*-values for these subsets are presented in green in **Figure S3c**.

We also implemented a clumping procedure on the 97,085,593 *k*-mers to quantify the overall amount of correlation in the entire dataset of 97,085,593 *k*-mers. The algorithm operated as follows: the *k*-mers list was sorted by sequence alphabetically. For a given threshold $0<\tau<1$, i) we selected the first *k*-mer and removed every *k*-mers such that $r^{2}\leq\tau$. We then moved to the next remaining *k*-mer and executed the same process until we reached the end of the list. We repeated this algorithm for $\tau$ in [0.6, 0.5, 0.4, 0.3, 0.2, 0.1, 0.05 and 0.01] and recorded the number of *k*-mers remaining after the filtering. The count of *k*-mers remaining is presented in **Figure S3d**.

### Supplementary Figures

#### Figure S1. Number of *k*-mers and genome length

Number of *k*-mer mapped to each species in the Milieu Interieur cohort as a function of the genome length. Both x and y axes are plotted on a log scale. Genome length was extracted from the NCBI database. When multiple lengths were available, the longest genome was used. The number of *k-*mers associated with a species was strongly proportional to its genome length (Pearson r = 0.77). On average, a species harboured 4 ,083 *k-*mers in the matrix (range = 1 – 7 ,938 ,905); 7 % of species had only a single *k-*mer. This is because the *k*-mers were strongly filtered, hence the genomes of low-abundance species are expected to be weakly covered. A subset of 77 species contained more than 1 million *k-*mers, the most extreme case being 7 ,938 ,905 *k-*mers.

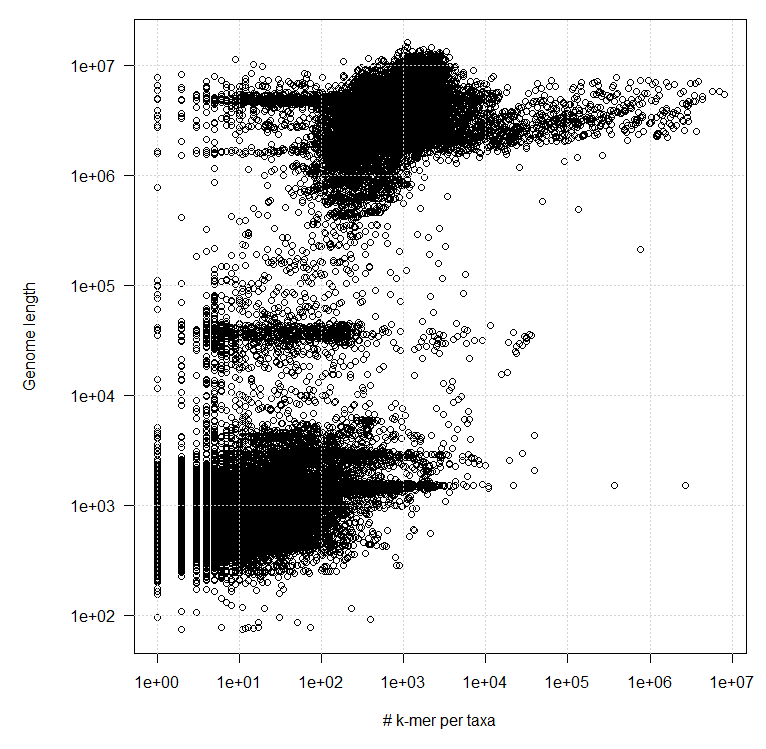

#### Figure S2. Species abundance from taxonomic classifiers and *k*-mers

Distribution of the correlation of species’ relative abundances between an estimate derived from the *k*-mer abundances and either Metaphlan4 (a) or Kraken2 (b) for a subset of 887 and 547 species quantified from both *k*-mer-based and Metaphlan4 or Kraken2. Correlations were derived for each of the 938 individuals from Milieu Interieur as cor(*k*-mer-based abundance, taxonomic classifier abundance).

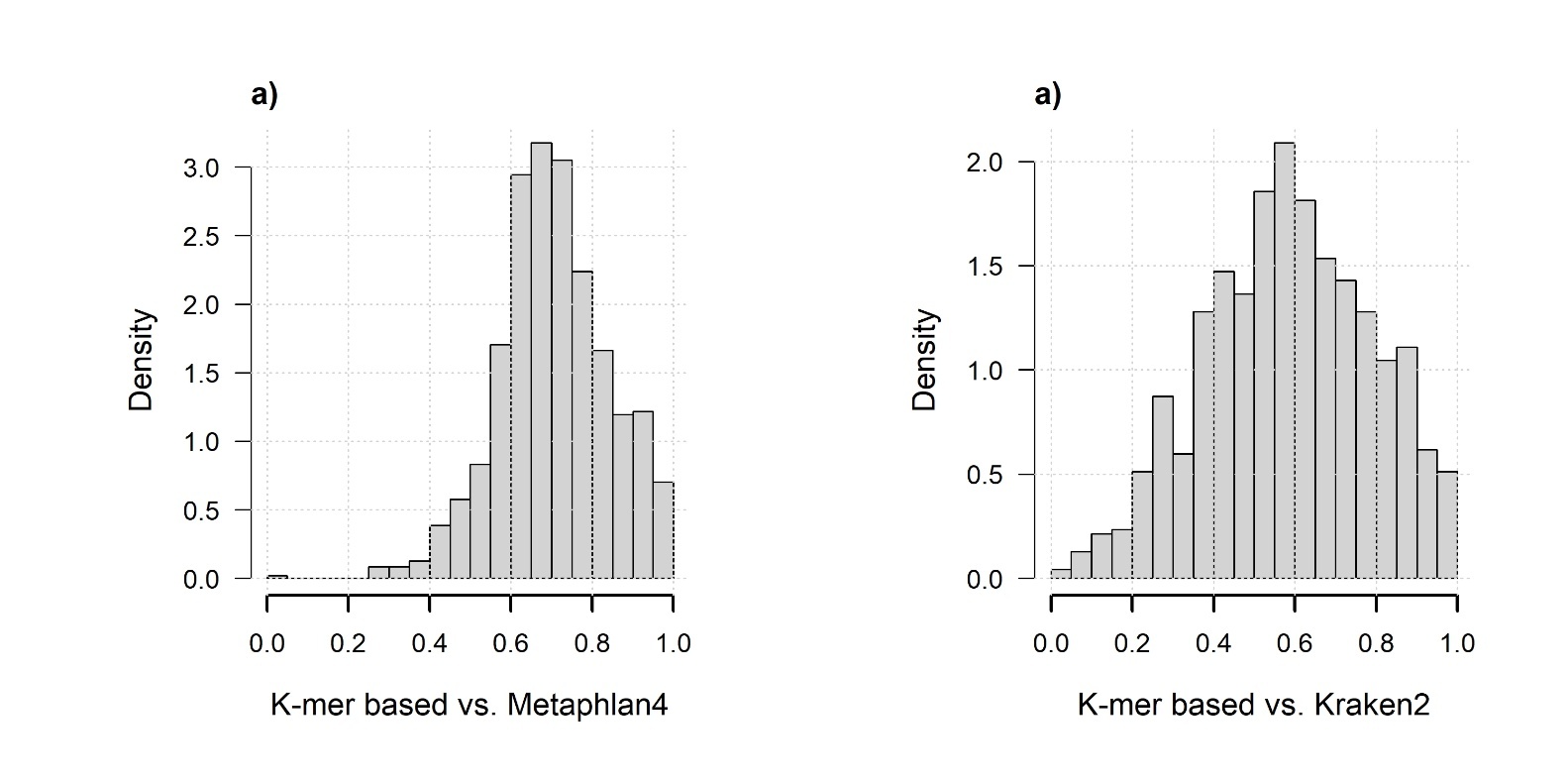

#### Figure S3. Calibration of MOGS under the null

We simulated 50 random phenotypes and conducted GWAS on the 97,085,593 *k-*mers derived from the Milieu Interieur. Panel a) presents the QQplot distribution, the observed -log10(P-value) as a function of the expected -log10(P-value). The red line and the red shade indicate the average and the 95% confidence interval across the 50 simulated phenotypes. The black line represents de $x=y$ axis. Panel b) presents the distribution across the 50 simulated phenotypes of the lambda value. For each simulated phenotype, the lambda is derived as the ratio of the median of the observed -log10(P) over the median of the expected -log10(P). Panel c) shows the QQplot distribution. The red line and the red shade indicate the average and the 95% confidence interval across the 50 simulated phenotypes. The green line and the green shade indicate the average and the 95% confidence interval across the 50 simulated phenotypes on 151,520 *k*-mers whose square correlation $r^{2}\geq0.5$. The blue line represents the average P-value distribution of 151,520 randomly selected *k*-mers across the 50 simulated phenotypes. The black line represents de $x=y$ axis. Panel d) provides the number of uncorrelated *k*-mers remaining after a clumping procedure for different square correlation threshold $r^{2}$.

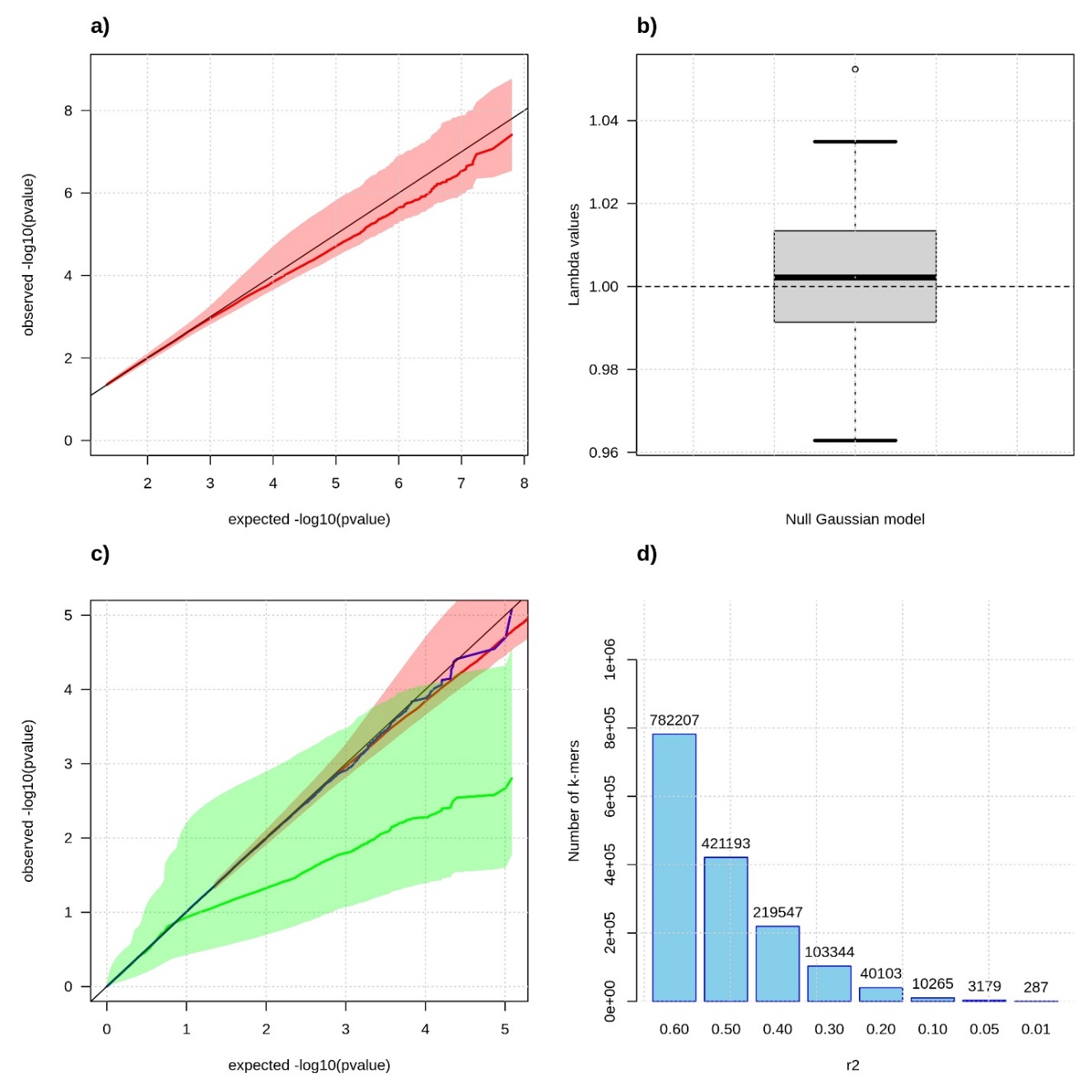

#### Figure S4. Phenotypic correlation in Milieu Interieur

Pairwise correlation between the 26 phenotypes from Milieu interieur covering demographics, physiological measurements, medical history, mental health, diet, physical activity and smoking habits.

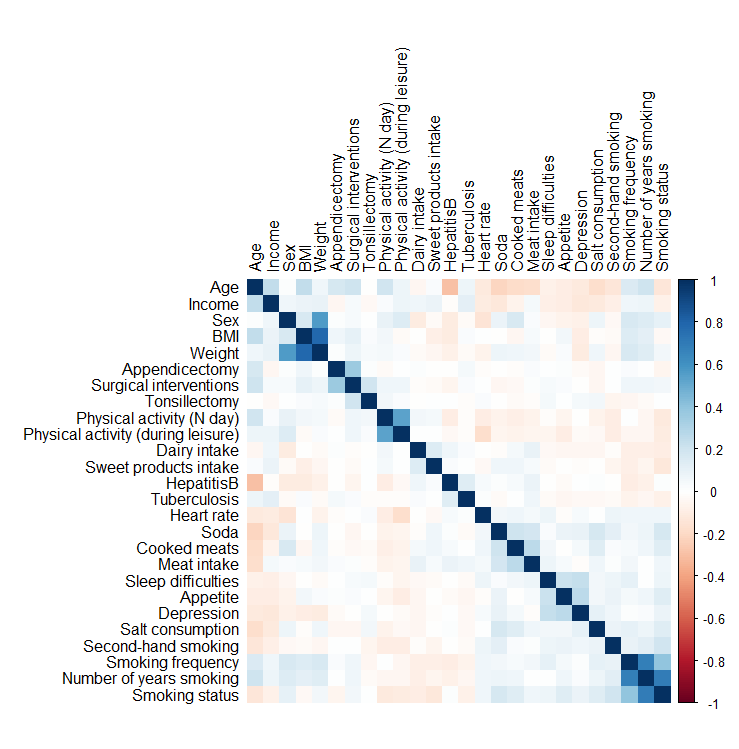

#### Figure S5. Genetic correlation at top associated *k*-mers

Pairwise correlation between the *k*-mer association statistics across the 93,905 *k*-mers associated with at least one phenotype (upper up triangle), against phenotype correlation (lower left triangle). This indicates multiple trends. For example, *k*-mers associated with increased age of the participant tend to be associated with lower smoking frequency, lower depression event, and lower appetite. *k*-mers associated with appetite are also positively associated with soda consumption and depression events.

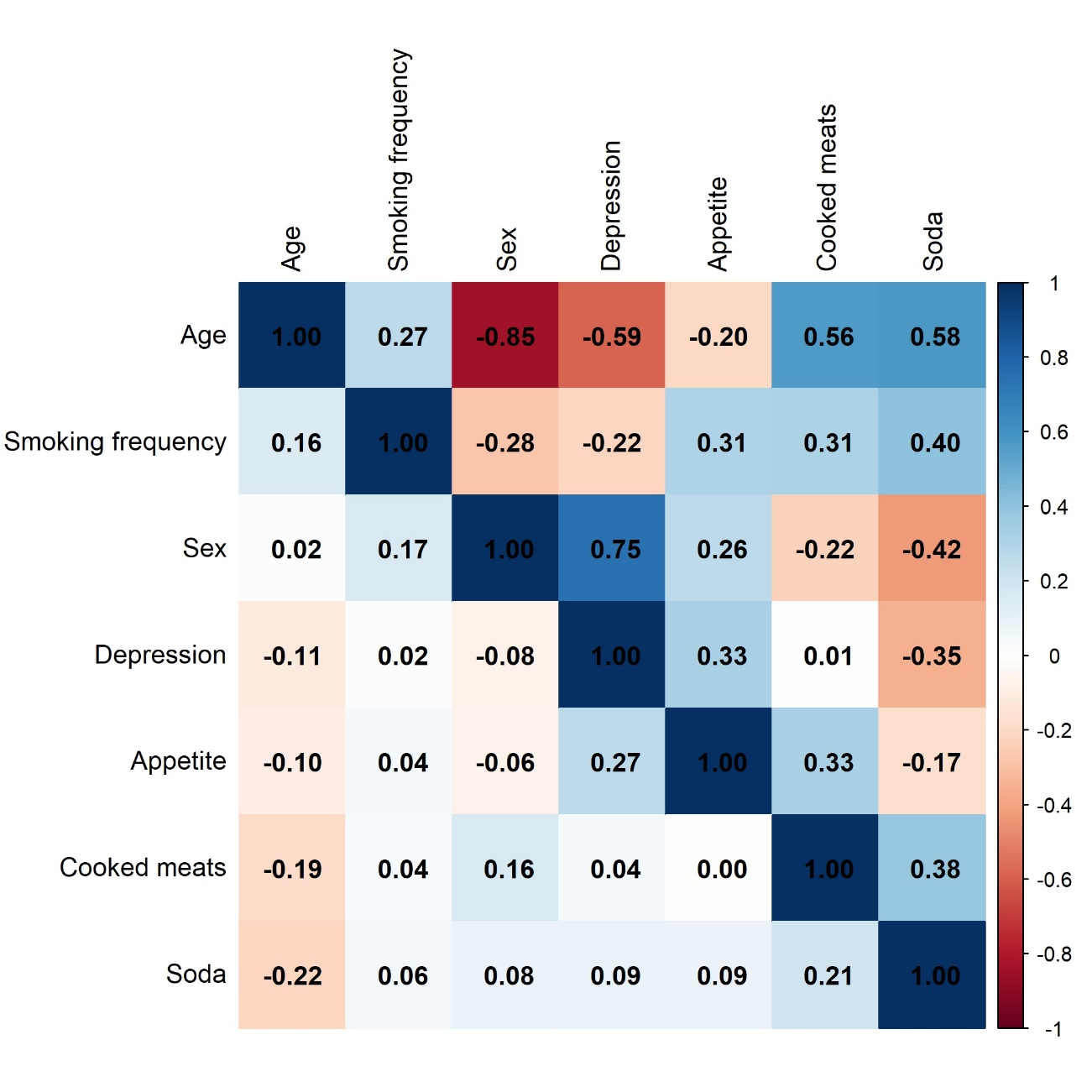

#### Figure S6. Correlation and annotation of *k*-mers associated with Age

Characteristics of the top 1,000 *k*-mers associated with *Age*. Panel a) shows the -log10(P-value) from all associated *k*-mers. Panel b) shows the correlation matrix between the *k*-mers and the associated dendrogram. *k*-mers were clustered using a ward.D2 algorithm applied to the correlation matrix. The position of the independent signal (IS) is presented below the correlation matrix. Panel c) presents the distribution of the annotations within each cluster. Annotations representing more than 5% of the total are labelled. *k*-mers unannotated or mapping to a “hypothetical protein” are indicated in grey.

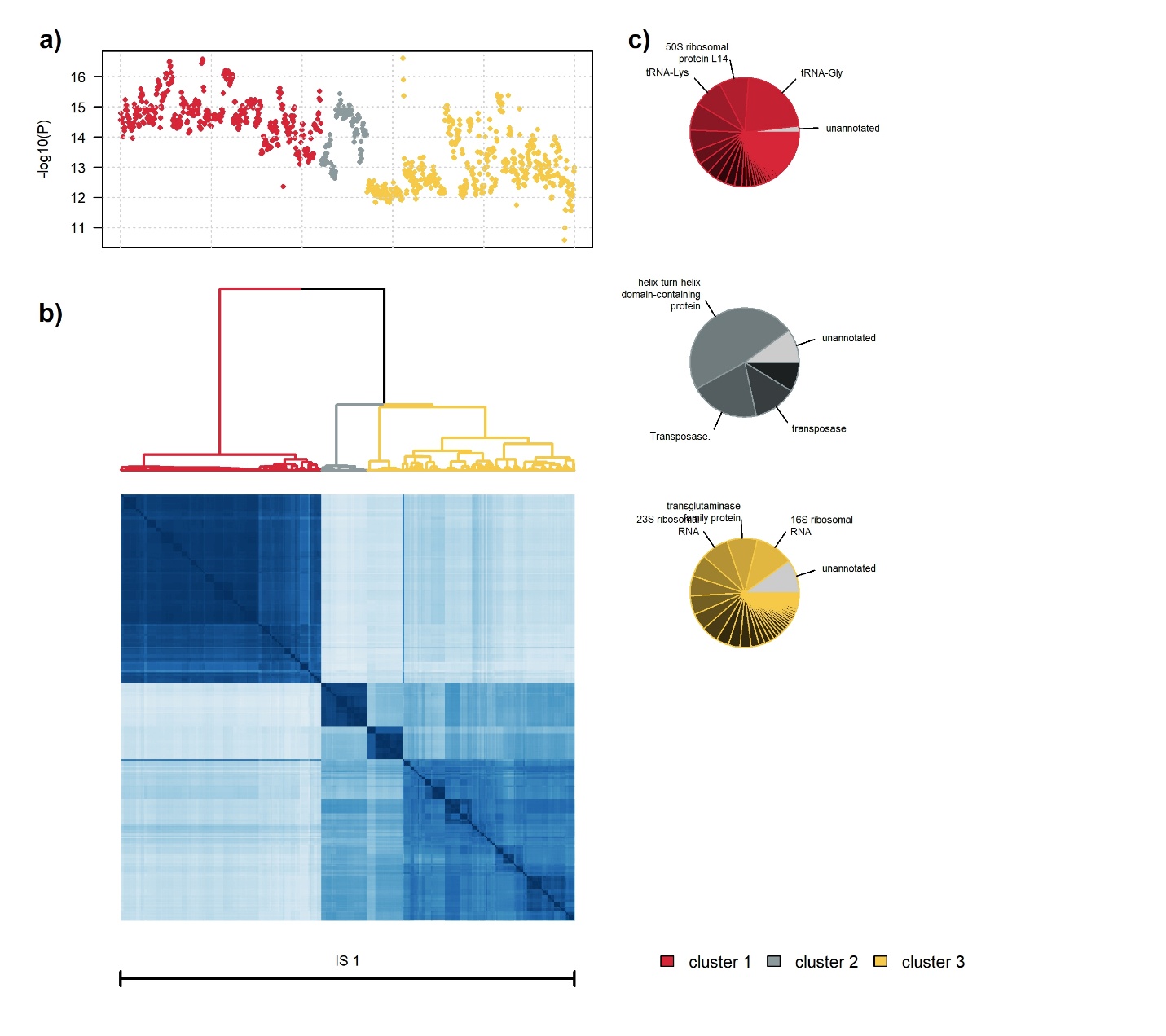

#### Figure S7. Correlation and annotation of *k*-mers associated with Appetite

Characteristics of the 393 *k*-mers associated with *Appetite*. Panel a) shows the -log10(P-value) from all associated *k*-mers. Panel b) shows the correlation matrix between the *k*-mers and the associated dendrogram. *k*-mers were clustered using a ward.D2 algorithm applied to the correlation matrix. The position of the independent signal is presented below the correlation matrix. Panel c) presents the distribution of the annotations within each cluster. Annotations representing more than 5% of the total are labelled. *k*-mers unannotated or mapping to a “hypothetical protein” are indicated in grey.

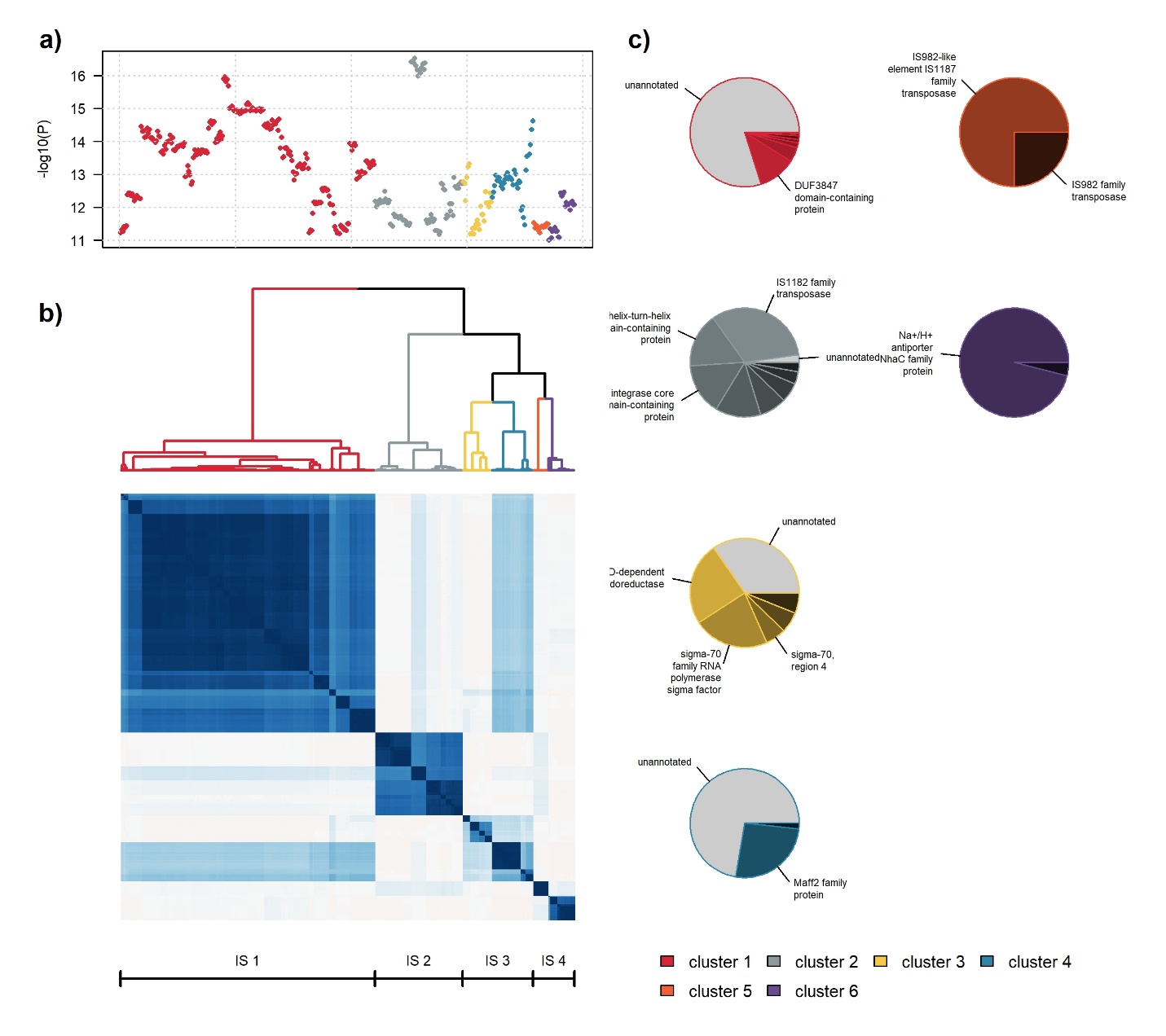

#### Figure S8. Correlation and annotation of *k*-mers associated with Depression

Characteristics of the 1,965 *k*-mers associated with *Depression*. Panel a) shows the -log10(P-value) from all associated *k*-mers. Panel b) shows the correlation matrix between the *k*-mers and the associated dendrogram. *k*-mers were clustered using a ward.D2 algorithm applied to the correlation matrix. The position of the independent signal is presented below the correlation matrix. Panel c) presents the distribution of the annotations within each cluster. Annotations representing more than 5% of the total are labelled. *k*-mers unannotated or mapping to a “hypothetical protein” are indicated in grey.

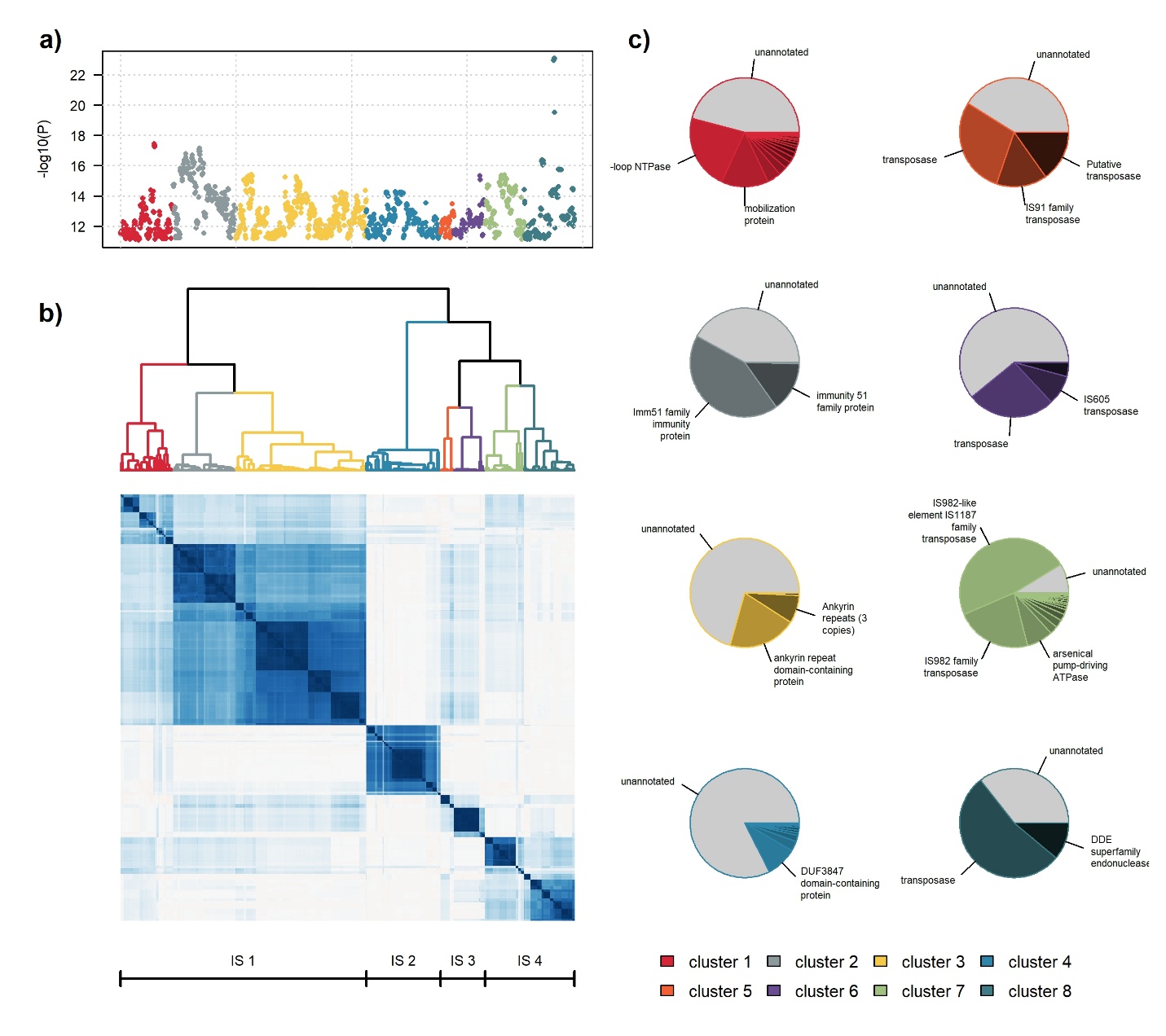

#### Figure S9. Correlation and annotation of *k*-mers associated with Cooked Meat

Characteristics of the 121 *k*-mers associated with *Cooked Meat*. Panel a) shows the -log10(P-value) from all associated *k*-mers. Panel b) shows the correlation matrix between the *k*-mers and the associated dendrogram. *k*-mers were clustered using a ward.D2 algorithm applied to the correlation matrix. The position of the independent signal is presented below the correlation matrix. Panel c) presents the distribution of the annotations within each cluster. Annotations representing more than 5% of the total are labelled. *k*-mers unannotated or mapping to a “hypothetical protein” are indicated in grey.

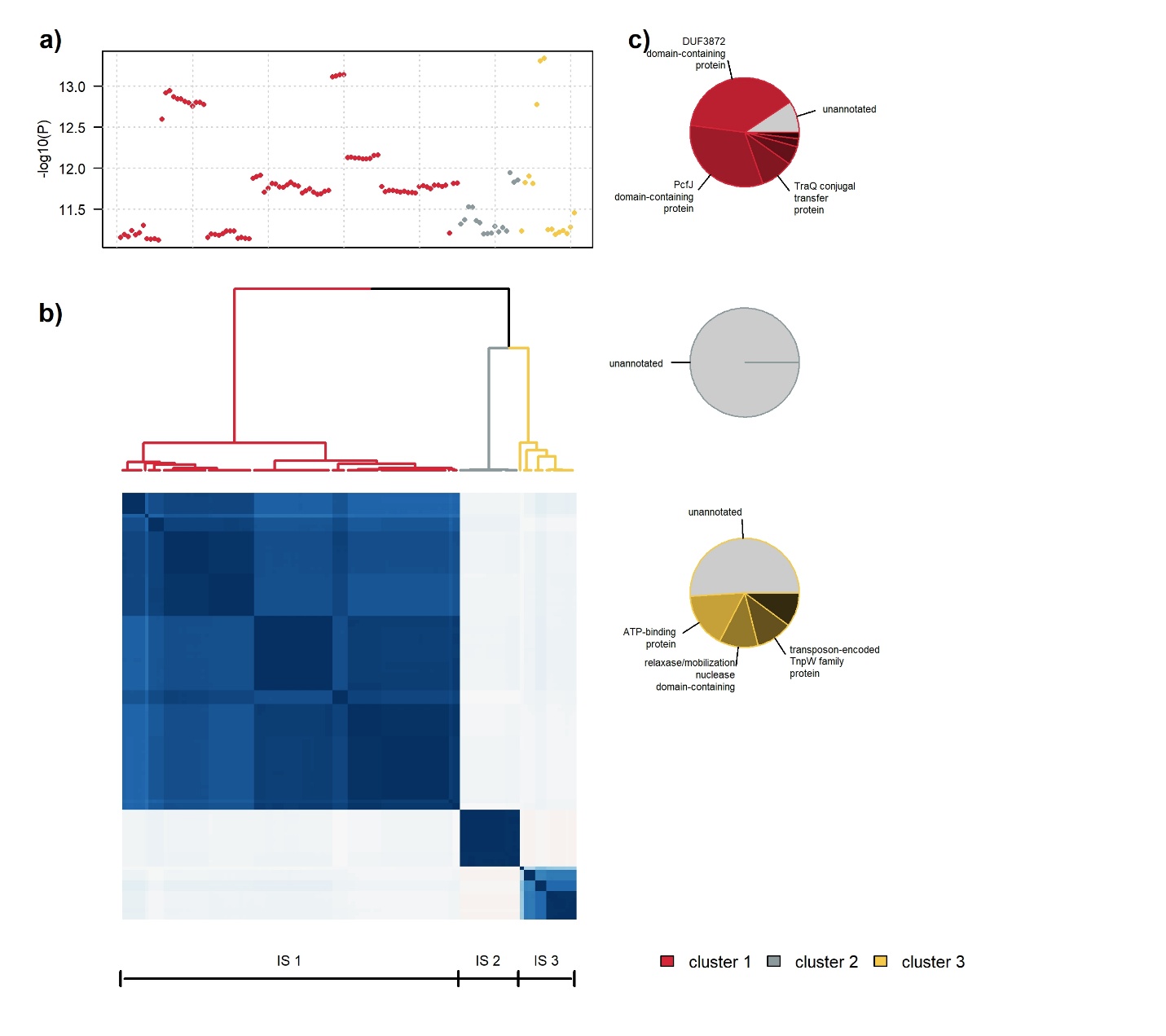

#### Figure S10. Correlation and annotation of *k*-mers associated with Soda

Characteristics of the 125 *k*-mers associated with *Soda consumption*. Panel a) shows the -log10(P-value) from all associated *k*-mers. Panel b) shows the correlation matrix between the *k*-mers and the associated dendrogram. *k*-mers were clustered using a ward.D2 algorithm applied to the correlation matrix. The position of the independent signal is presented below the correlation matrix. Panel c) presents the distribution of the annotations within each cluster. Annotations representing more than 5% of the total are labelled. *k*-mers unannotated or mapping to a “hypothetical protein” are indicated in grey.

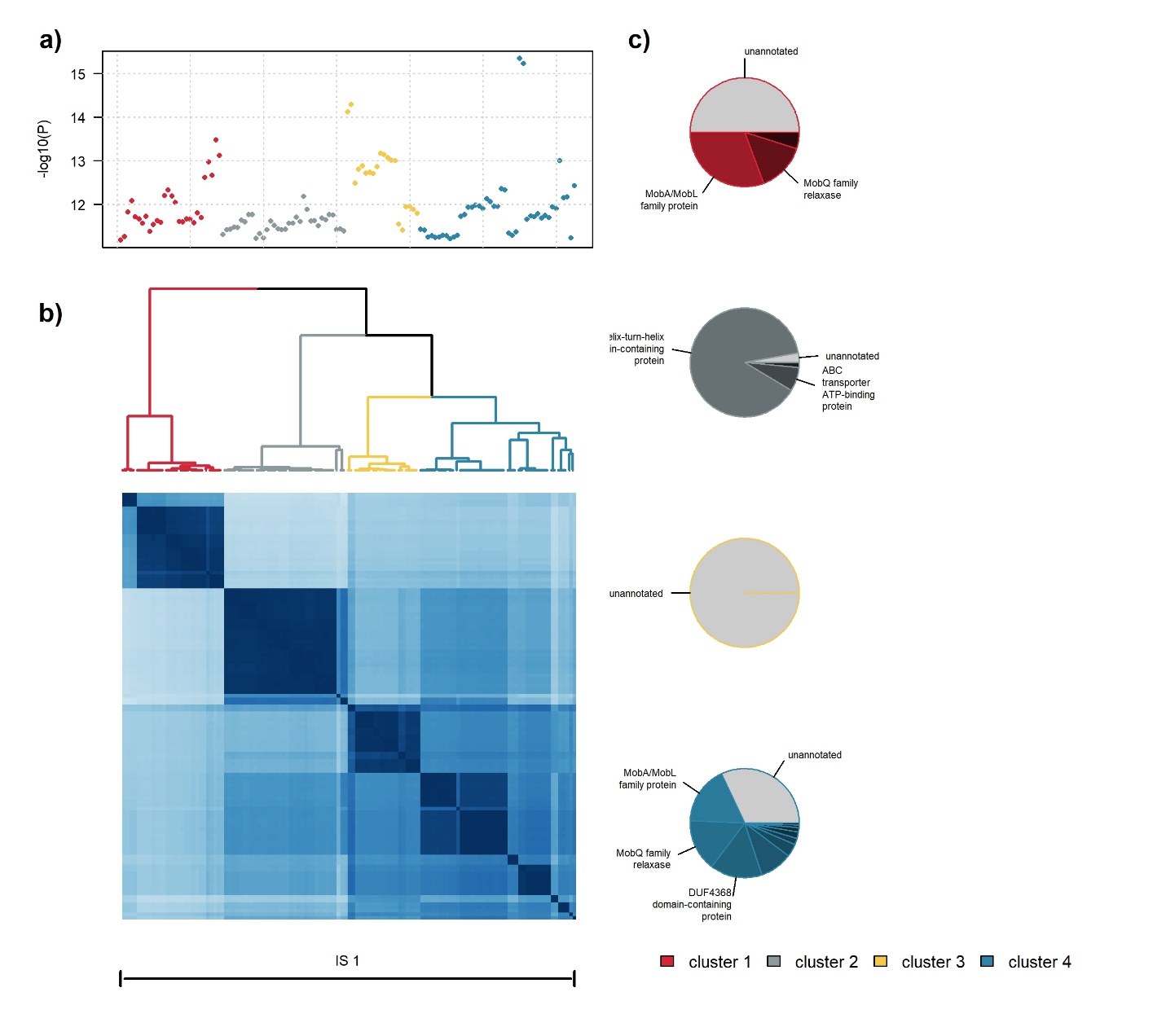

#### Figure S11. Correlation and annotation of *k*-mers associated with Sex

Characteristics of the 57 *k*-mers associated with *Sex*. Panel a) shows the -log10(P-value) from all associated *k*-mers. Panel b) shows the correlation matrix between the *k*-mers and the associated dendrogram. *k*-mers were clustered using a ward.D2 algorithm applied to the correlation matrix. The position of the independent signal is presented below the correlation matrix. Panel c) presents the distribution of the annotations within each cluster. Annotations representing more than 5% of the total are labelled. *k*-mers unannotated or mapping to a “hypothetical protein” are indicated in grey.

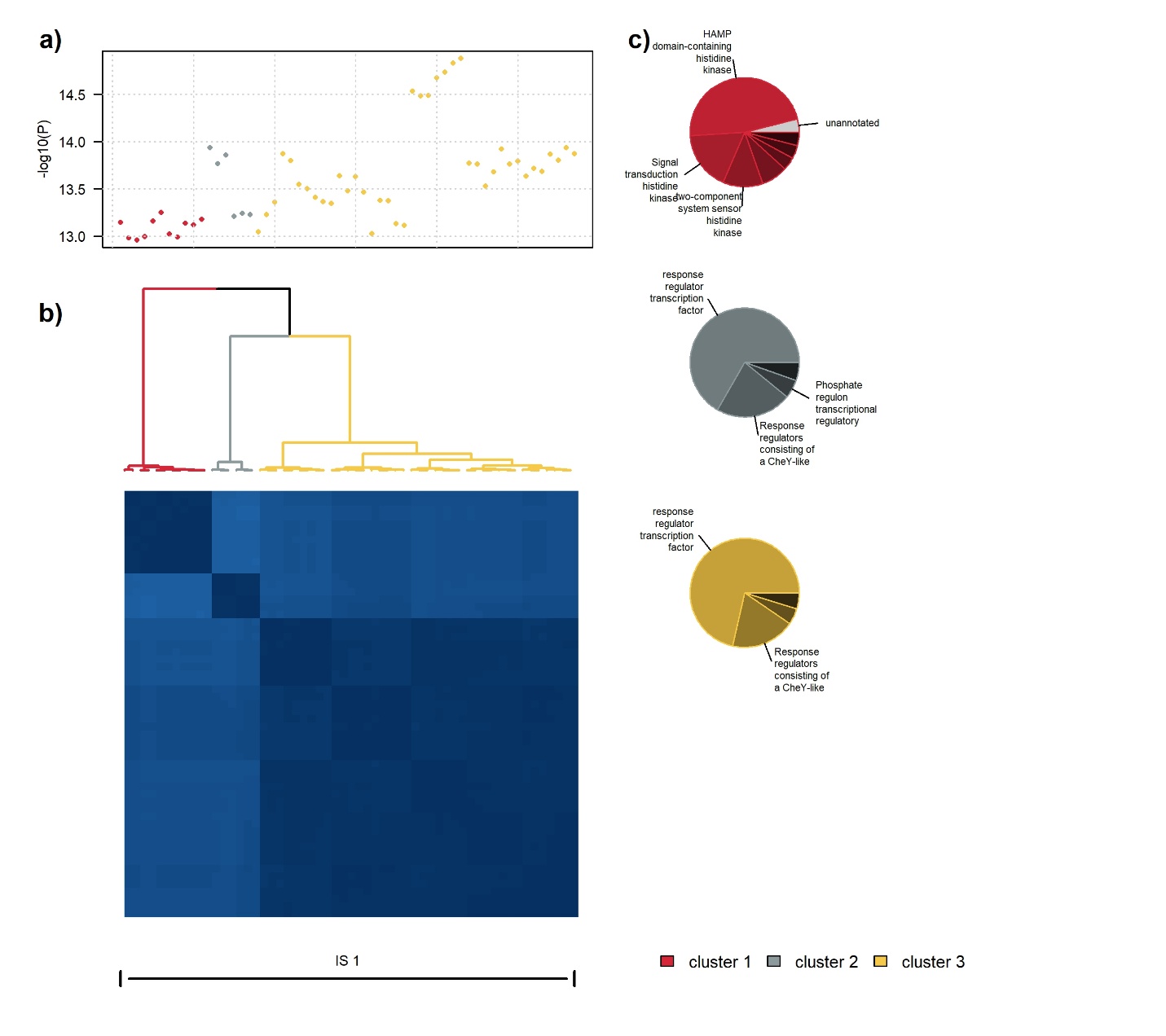

#### Figure S12. Correlation and annotation of *k*-mers associated with Smoking Frequency

Characteristics of the 5,310 *k*-mers associated with *Smoking Frequency*. Panel a) shows the -log10(P-value) from all associated *k*-mers. Panel b) shows the correlation matrix between the *k*-mers and the associated dendrogram. *k*-mers were clustered using a ward.D2 algorithm applied to the correlation matrix. The position of the independent signal is presented below the correlation matrix. Panel c) presents the distribution of the annotations within each cluster. Annotations representing more than 5% of the total are labelled. *k*-mers unannotated or mapping to a “hypothetical protein” are indicated in grey.

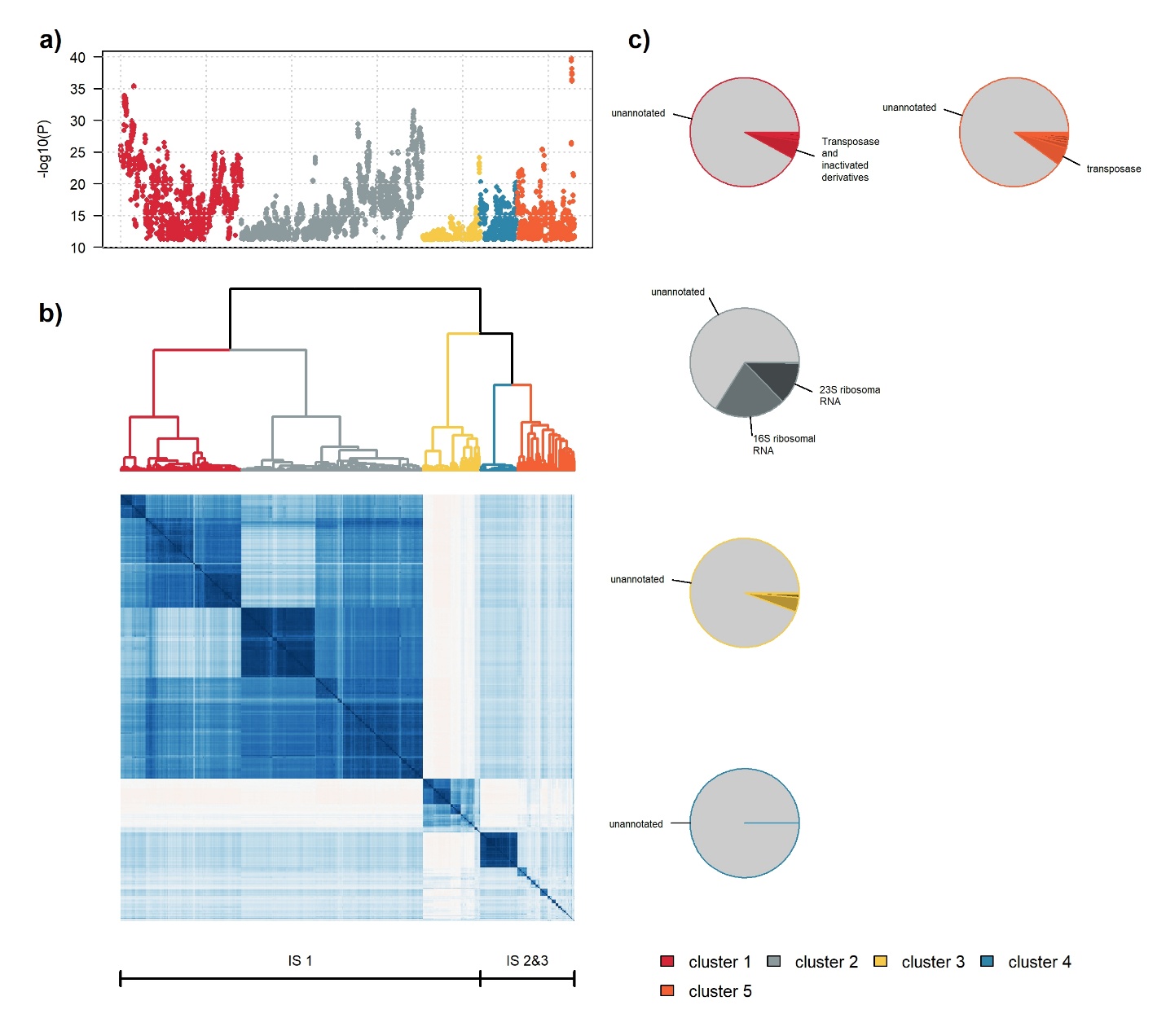

#### Figure S13. Species-based screening analysis in Milieu Interieur

We derived genes for the 938 individuals from the Milieu Interieur cohort using *HUMAnN*. Panel (a) presents and compares two distributions of the mean relative abundance in reads per kilobase (RPK) per genes across the individuals. Distribution in red includes all the 118,628 genes found by *HUMAnN* while distribution in blue covers only the 619 genes significantly associated with age phenotype. Panel (b) presents the gene-phenotype association QQplot of all the phenotypes highlighting the seven phenotypes reaching a stringent Bonferroni corrected significance threshold (horizontal red dash line, P < 1.62 x 10^-8^).

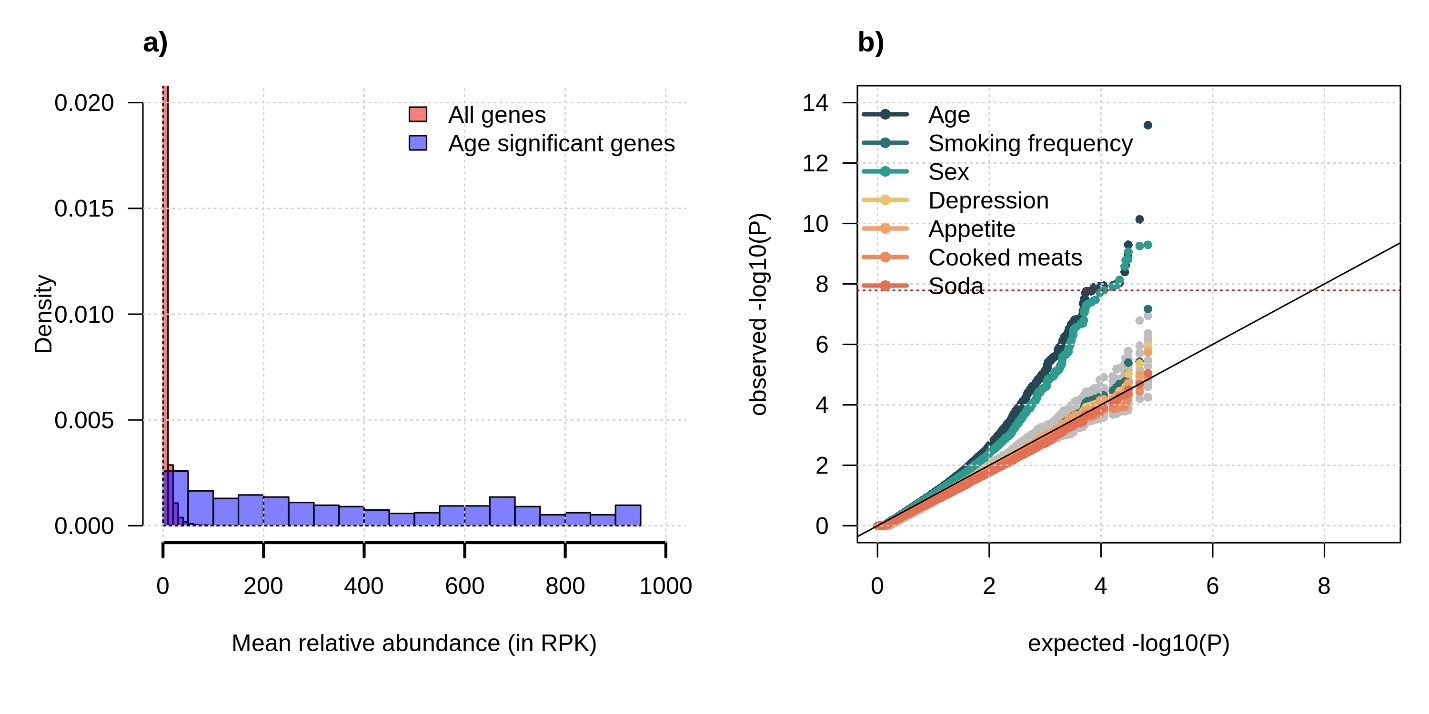

#### Figure S14. Evaluation of MOGS software runtime by sample size and *k*-mer count

Benchmark of MOGS software runtime for different sample sizes and numbers of *k*-mers. Experiments were based on simulations. The mean time (in seconds) was estimated on 3 executions of MOGS.

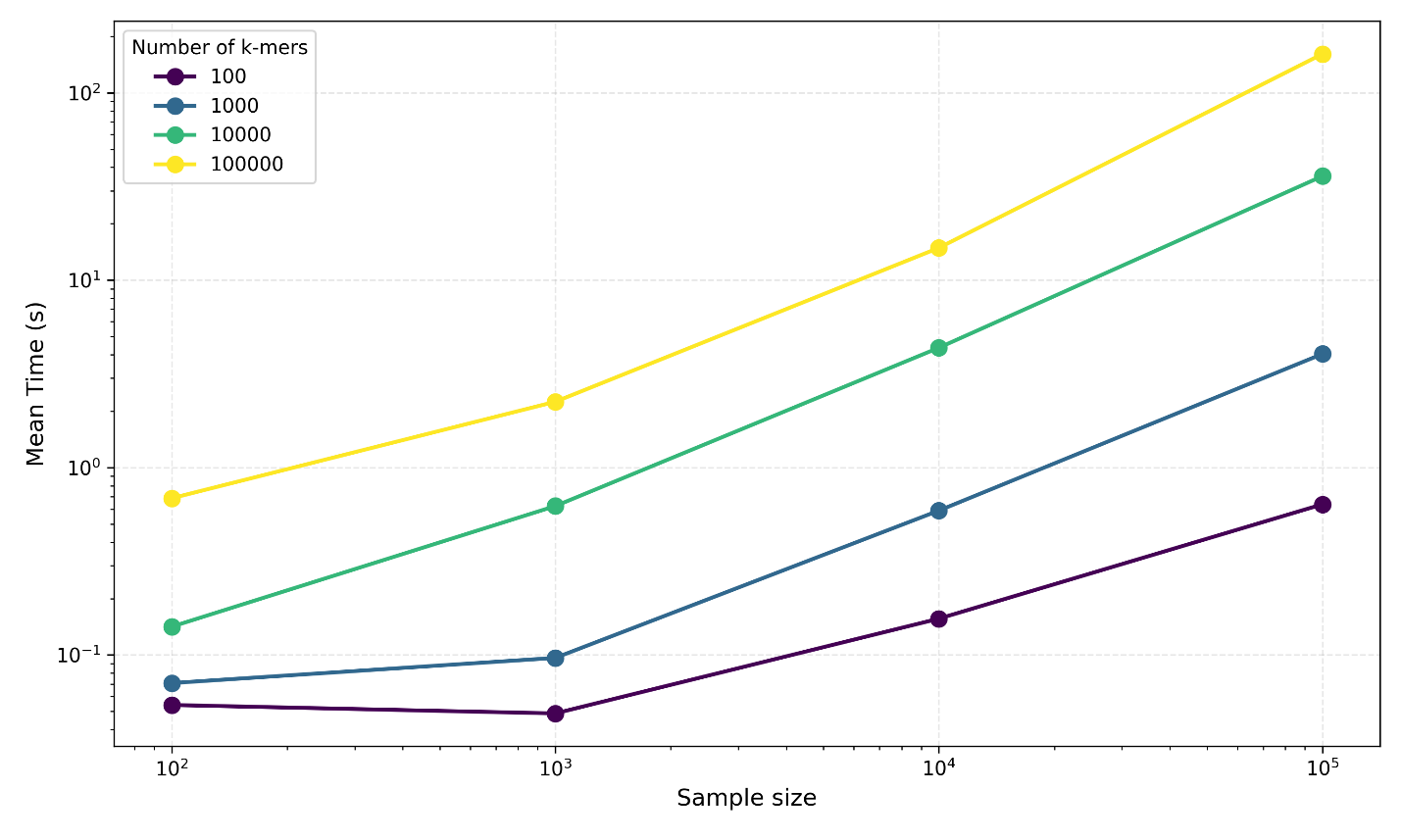
